# The importance of DNA sequence for nucleosome positioning in transcriptional regulation

**DOI:** 10.1101/2023.08.01.550795

**Authors:** Malte Sahrhage, Niels Benjamin Paul, Tim Beißbarth, Martin Haubrock

## Abstract

Nucleosome positioning is a key factor for transcriptional regulation. Nucleosomes regulate the dynamic accessibility of chromatin and interact with the transcription machinery at every stage. Influences to steer nucleosome positioning are diverse, and the according importance of the DNA sequence in contrast to active chromatin remodeling has been subject of long discussion. In this study, we evaluate the functional role of DNA sequence for all major elements along the process of transcription. We developed a random forest classifier based on local DNA structure that assesses the sequence-intrinsic support for nucleosome positioning. On this basis, we created a simple data resource that we applied genome-wide to the human genome. In our comprehensive analysis, we found a special role of DNA in mediating the competition of nucleosomes with cis-regulatory elements, in enabling steady transcription, for positioning of stable nucleosomes in exons and for repelling nucleosomes during transcription termination. In contrast, we relate these findings to concurrent processes that generate strongly positioned nucleosomes in vivo that are not mediated by sequence, such as energy-dependent remodeling of chromatin.

**GRAPHICAL ABSTRACT:** 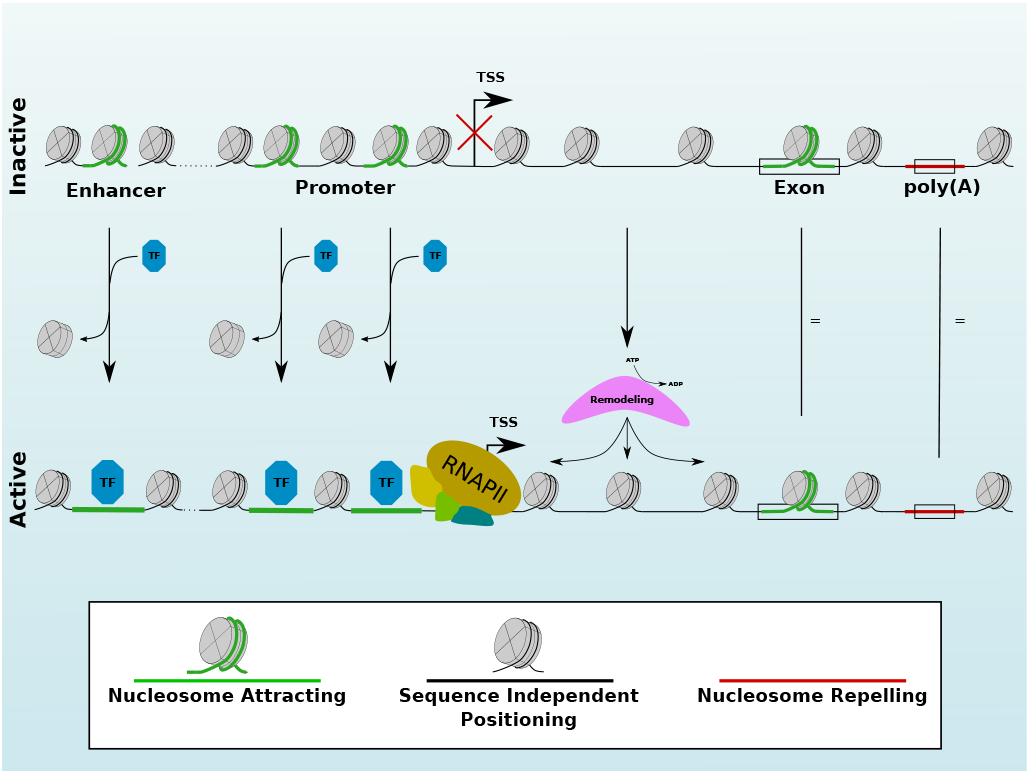

## INTRODUCTION

The nucleosome is a foundational protein complex in eukaryotic organisms formed by an octamer of pairs of histone-subunits (two pairs of H2A/H2B and a tetramer of H3/H4). This complex wraps 147 base-pairs (bp) of DNA (1, 2). The positioning of nucleosomes is very important for the regulation of transcription and the relevance of nucleosome positioning for the regulation of genes can be viewed from many perspectives. The broadest unit to span interaction potential are topologically associated domains (TADs). These are characterized by a loop compartment that brings together potentially interacting proteins at enhancers and promoters of genes close to each other (3). The boundaries of these logical units are set up at so called insulator regions which are defined by an interplay of CTCF binding (4) and a consequential strict positioning of a well-positioned nucleosome array (5). Whether the genes inside such units are transcribed is an intricately regulated process involving nucleosome positioning at all levels. The binding of transcription factors (TFs) to promoter and enhancer regions is crucial for the initiation of transcription. As a means of controlling access to transcription factor binding sites (TFBSs), nucleosomes occupy DNA and need active displacement through (pioneer-)TFs or energy dependent remodeling to avoid random triggering of gene expression (6, 7). To achieve an orderly rate of transcription, the RNA polymerase II (RNAPII) is highly regulated by nucleosomes. In particular, the +1 nucleosome directly downstream of the transcription start site (TSS) and a sequential array of well-positioned nucleosomes is needed to reliably initiate transcription (8, 9, 10). Along its way, RNAPII encounters more fuzzily positioned nucleosomes in intronic regions and well-positioned nucleosomes in exons. Since exonic DNA is the template for the subsequent translation of proteins, the positioning of nucleosomes supports the faultless transcription of these regions by serving as roadblocks to decrease the speed of RNAPII elongation, by marking alternative splice sites (11) and by triggering feedback signals for other concurrent transcription initiation events on the same gene (12). The transcription ends at the transcription termination site (TTS), which sets up the signal where polyadenylation will be applied on the resulting pre-mRNA. These polyadenylation sites (poly(A) sites) are known to exhibit a considerate nucleosome depletion directly around the DNA encoded prospective polyadenylation site compared to their nucleosome occupied surroundings (13). An extra layer of nucleosome functionality in all of these stages is added through the epigenetic modifications of histone proteins. Depending on the specific biochemical modulations of the histone sub-units, nucleosomes can carry marks for the active recruitment of further proteins and show a different flexibility in terms of the energy needed for displacement (14, 15). Since the differential increase or decrease of chromatin accessibility at all of these specific functional sites regulates transcription and thus cell identity, it is crucial to understand the principles that determine nucleosome positioning.

The influences on the positioning of nucleosomes have been debated for many years and can be grouped roughly into DNA sequence, trans-acting factors and active chromatin remodeling (16, 17, 18). The role of the DNA sequence is particularly controversial. It was postulated that there is a periodic repeat of A/T di-nucleotides that support the binding of nucleosomes (19, 20, 21). This pattern has been extended to be represented by local DNA structure (22) with the help of DNAshape (23). The benefits of the supposed patterns for nucleosome support are linked to the flexibility of the DNA strand to facilitate an easier attraction of nucleosomes (24, 25), which can be seen particularly well in extremely stiff poly(dA:dT) tracts that counteract nucleosome binding (26).

Some of these findings have been discovered with the help of a variety of computational tools to predict nucleosome positions, each with a different focus of training data, prediction target and methodology (27, 28, 29, 30). Among these, there are tools that employ HMMs (31, 32), SVMs (33) and recently more and more CNN architectures (34, 35, 36, 37). The difference in meaning between sequence-intrinsic signals and the actual *in vivo* positioning of nucleosomes is fundamental to understand. The sequence can only explain the potential of any given position to attract nucleosomes. Whether it is actually occupied by nucleosomes or opened up by regulatory mechanisms cannot be determined without the help of experimental chromatin accessibility data for the particular cell type. In other words, while experimental data can provide insights into the actual nucleosome positioning within cells, it does not directly elucidate the underlying reasons or mechanisms behind such positioning. On the other hand, machine learning on DNA sequence can give an intrinsic estimate of nucleosome attraction potential but cannot answer the question where nucleosomes are located *in vivo*. Therefore, we need to integrate both information to estimate the overlap of sequence-intrinsic nucleosome support and experimental chromatin accessibility evaluation (38). For that purpose we use nucleosome occupancy data (MNase-seq) (39), chromatin accessibility data (DNase-seq) (40) and ChIP-seq data for both histone modifications and TFs, mainly in the two cell lines GM12878 and K562 available from ENCODE (41).

In this study, we evaluate the importance of DNA sequence for nucleosome positioning at all elements involved in transcription in the human genome. For that purpose, we developed the nucleosome formation score (NF score), which describes the sequence-intrinsic support for attracting nucleosomes purely based on DNA. The score is derived using a classification algorithm which is applied in a genome-wide manner. We provide it as a simple data resource which can be used without further computation for similar analyses in the future. Due to its relative robustness against overfitting we created a random forest (42) classifier based on the Guo2014 dataset (33) of nucleosomal sequences which is derived from MNase-seq in human CD4^+^ T-cells (39). Our classifier uses a transformation from raw DNA sequences to a frequency spectrum of local DNAshape (Propeller Twist, Helix Twist, Minor Groove Width, Electrostatic Potential, obtained with the DNAshapeR package (43)). This binary classifier was applied in a sliding-window manner to the human genome in two different resolutions, yielding a genome-wide mapping of sequence-intrinsic nucleosome support.

We aspire to give a comprehensive overview over the crucial parts of the transcriptional machinery and provide our genome-wide measure for sequence-intrinsic nucleosome support as a simple data resource. Our findings are embedded into a framework of previous studies that showed the relevance of DNA for nucleosome positioning for some specific elements and we add some new insights about the sequence-definition of nucleosome positioning in transcription. We demonstrate that there is a clear influence of DNA sequence for supporting nucleosome binding at positions of high competition with other DNA-binding proteins and where it is beneficial to accumulate nucleosomes by default, such as exons, or repel them in transcription termination sites. In contrast, we give examples of positioned, but sequence-independent nucleosome arrays in proximity to the TSS or around insulator sites and put our findings into the context of more unspecific, fuzzy nucleosomes.

## METHODS

### Datasets

The experimental chromatin accessibility datasets were obtained from ENCODE for the human genome in hg19 (41). They comprise of MNase-seq data, as density graph of signal enrichment described in (44), DNase-seq data as read depth normalized signal, described in (45), peak-called histone modification ChIP-seq data as bed narrowPeak locations for K562 and GM12878 and raw signal for H3K4me3 and H3K27ac as control normalized tag-density, described in (46). Besides, we used CTCF ChIP-seq peaks from ENCODE for our insulator analysis. Furthermore, we used the comprehensive list of ChIP-seq narrowPeak locations for all TFs that are available on ENCODE for hg19. When referring to the central peak position related to the ENCODE data, we reference the point-source peak defined in the narrowPeak format standard. Due to their extensive length, we show the ENCODE IDs for all the aforementioned datasets in Supplementary Table S1, Supplementary Table S2 and Supplementary Table S3.

Additionally, we used nucleosome occupancy data to evaluate the influence of the GATA3 TF in MDA-MB-23 cells with and without GATA3 expression. The according MNase-seq data was taken from (47). We calculated the mean of all three replicates for the condition of GATA3 expression and for the control cells without GATA3 expression.

Furthermore, we evaluated the evolutionary conservation at base pair-resolution with the PhyloP score (48), which was obtained using the UCSC table browser (49).

We compared all these data along a variety of different genome locations. The broad overview over different biological contexts is defined by chromHMM for K562 and GM12878 and defines 15 meaningful states originally derived from histone modification ChIP-seq data (50).

The gene mapping we use is based on RefSeq in hg19 (51) and was also obtained with the UCSC table browser. We exclude genes from the chromosome Y, since not all experimental chromatin accessibility data is available for it. In particular, we use the remaining 30142 unique TSS locations to define genes and promoters. Our promoter definition is defined as within 1 kb upstream of each TSS defined by RefSeq. In case of multiple alternative TSS locations, all of these were considered as individual gene starts. Besides using RefSeq for TSS, we also extracted all exon locations from the same mapping.

Moreover, we analyzed the elements at TTSs in the form of poly(A) sites. The poly(A) site mapping with a separation into clusters of biological origin is taken from PolyASite 2.0 (52). To obtain single locations for the center of each site, we used the representative cluster location from the unique cluster ID column. The original assembly of the data is hg38. We used liftOver to convert the coordinates to hg19 via the rtracklayer R package (53).

### Software

Apart form the individual software that is referenced in its due place, the analyses were performed using R (v. 4.3) (54) and we used the packages rtracklayer (v. 1.60.0) (53), ggplot2 (v. 3.4.2) (55), GenomicRanges (v. 1.52.0) (56), ROCR (v.1.0.11) (57) and randomForest (v. 4.7.1.1) (58).

### Classifier

The classifier is based on the human Guo et al. 2014 dataset (33) which is in turn derived from an *in vivo* mapping of nucleosome positions in CD4^+^ T-cells (39). It contains 4573 non-redundant FASTA sequences which are categorized as either nucleosomal (n=2273) or linker sequences (n=2300). The FASTA files are transformed with the DNAshapeR tool (v. 1.28.0) (43), resulting in a set of numerical vectors for Roll, HelT, ProT and MGW with one value per bp each. Additionally, a power spectrum (Power Spectral Density, PSD) is applied to the DNAshape values in order to strengthen positional independence of the periodic nucleotide patterns, emphasizing the assumed underlying periodic sequence pattern and making it suitable for the use in a random forest classifier. That step uses the spectrum function in R with a taper of 0.1 and a span of 10. In a next step, the resulting spectral densities are concatenated into a feature matrix which is the input for a random forest classifier. The random Forest classifier was trained with the randomForest package in R with default parameters. The result of the classification is the categorization of a sequence being either *nucleosomal* or *linker* and serves as the basis for the two types of NF scores that will be described further in the NF score section.

For a better understanding about how clearly the data can be expected to be separated, Figure 1 C shows a t-distributed stochastic neighbor embedding (t-SNE) representation of the data used to train the classifier. The t-SNE method allows for visualizing high-dimensional data in two dimensions. In contrast to the common method of dimension reduction with principal component analysis (PCA), this method is able to capture non-linear relations of hidden data dimensions. It can be seen that the training data does not separate into two clearly distinct clusters. However, there is a decent separation between the nucleosomal training sequences (red) and the linker sequences (black), indicating a decent chance to categorize both classes, but no perfect prediction accuracy is to be expected with any method on that dataset. The performance of the classifier has a class-weighted accuracy of 0.815, evaluated with 10-fold cross-validation, and an area under ROC curve (AUC) of 0.883. The ROC curve for the classifier can be seen in Figure 1 B. The most recent classifiers use convolutional neural networks (CNNs) and reach accuracies of up to 0.889 (CoreNUP (59)) and an AUC of up to 0.94 (LeNUP (35)) on the given dataset. However, although CNNs are undoubtedly valuable for such prediction tasks in principle, we opted not to use them on this specific dataset due to their propensity for overfitting on smaller datasets.

**Figure 1.**
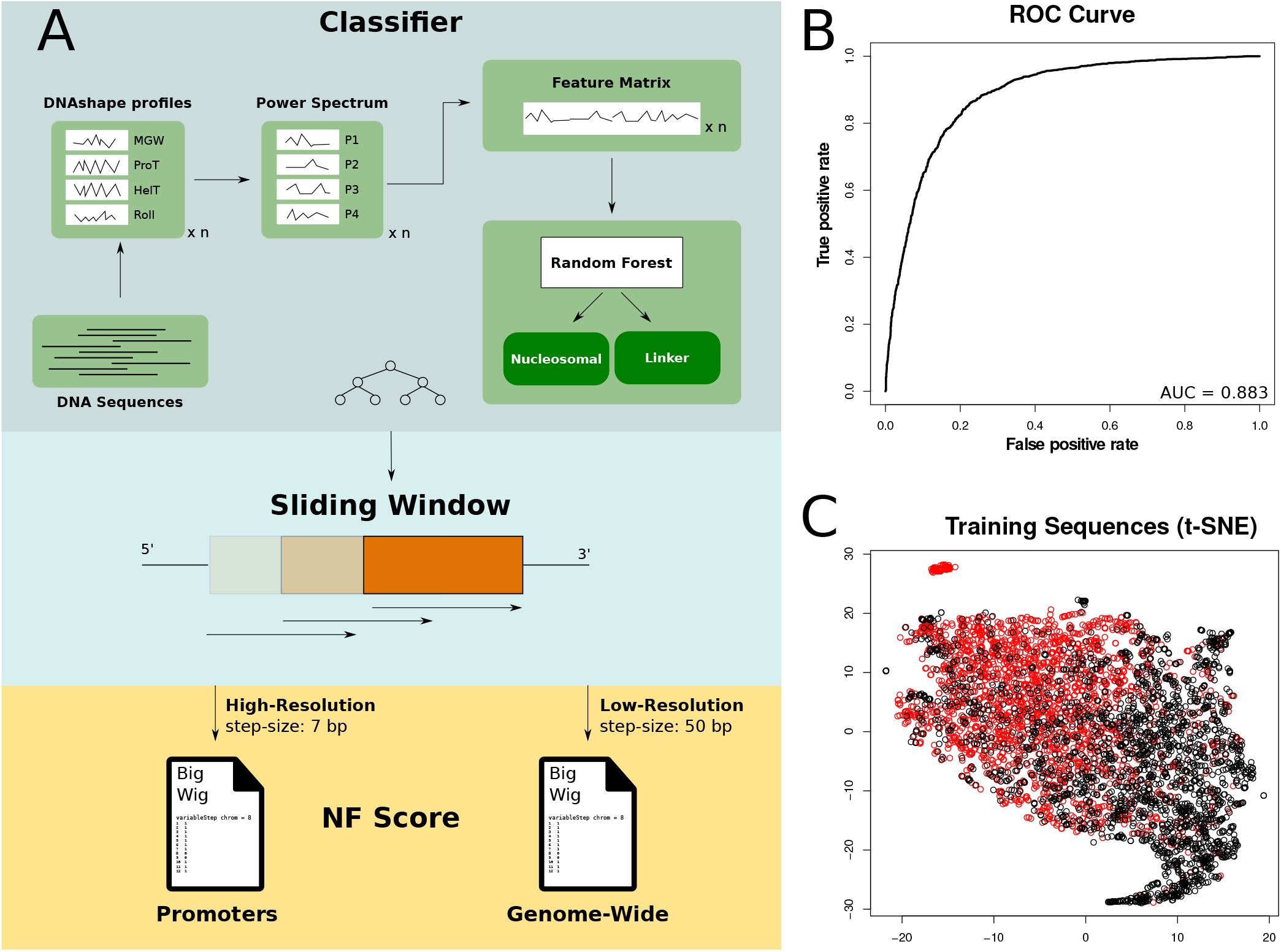
Processing and Classification of DNA Sequences. **(A)** Deriving the NF score. A random forest model is used to annotate DNA sequences as nucleosomal or linker. To build the feature matrix, each DNA fragment of length 147 bp is transformed into a frequency spectrum of local DNAshape. In order to create a pre-calculated mapping for the human genome, we applied the classifier genome-wide in a sliding-window approach with a step size of 50 bp and a detailed version for promoters with a step size of 7 bp. Both resources are provided as a downloadable bigWig file (see data availability section). **(B)** Receiver Operating Characteristic (ROC) curve and area under ROC curve (AUC). **(C)** t-SNE representation of training data based on the local DNAshape frequency spectrum to show how well the underlying data is separable in principle. Red dots represent nucleosomal and black dots indicate linker sequences. The non-linear dimension reduction shows that the two groups are overall separable in principal but there is no simple line to distinguish between both datasets. Thus, an upper boundary for the classification performance is to be expected based on the chosen training data.

The present *in vivo* training dataset is a common standard in the prediction of nucleosome positioning. We want to point out that the utilization of *in vivo* data has the potential to introduce bias in predicting nucleosome positioning events, independent of the chosen classification method. While previous studies have demonstrated the promise of incorporating *in vitro* nucleosome maps (60, 61), we find it noteworthy that despite training on *in vivo* data, we can show that the classifier is identifying the underlying sequence importance rather than the *in vivo* nucleosome occupancy that is resulting from the dynamic displacement of sequence-positioned nucleosomes in living cells. This is especially evident in the chapters about the counteracting relationship of high sequence-intrinsic nucleosome support in regulatory important regions against the low actual nucleosome occupancy *in vivo* at these positions of nucleosome-TF competition.

### Score Calculation

#### Nucleosome Formation (NF) score

The classifier described above gives a binary prediction for a DNA segment of 147 bp in length. To allow for an easier handling of the estimates of nucleosome support based on nucleotide sequence, we pre-calculated two sets of scores for the human genome in hg19 assembly with different local resolutions. (1) The low-resolution score is a simple application of the classifier in a sliding window to all regions of the human genome with a fragment input length of 147 bp and a step size of 50 bp. The resulting NF score is an estimate of either 1 (nucleosomal) or 0 (linker) for the central 50 bp in any given 147 bp bin. (2) The high-resolution NF score provides a continuous score between 0 and 1 and is based on a much more local sliding window with a segment size 147 bp and a step size of 7 bp. All base-pairs are thus part of multiple (147/7=21) predictions and the score represents the relative frequency of nucleosomal prediction results, this particular bp has been part of. To give an example, if an individual bp has been part of 21 sliding windows, of which 14 were classified as nucleosomal and 7 as linker, then the score at that base-pair is 14/21= 0.67. Due to the introduction of artifacts otherwise the step size needs to be a divisor of the window size of 147 bp. Thus we chose a resolution of 7 bp as it is the most local one next to a resolution of 1 bp.

Due to resource efficiency the high-resolution score was only pre-calculated for all human promoter regions, since we want to give a good local estimate of the importance of the sequence for nucleosome positioning in specific individual promoter regions. In contrast to the high-resolution score, the step size of 50 for the low-resolution score is purely pragmatical since there is no further integration of the classification results. If not specified otherwise, we always use the low-resolution score, since we normally show the mean of nucleosome support profiles over a large number of fragments and there is virtually no difference between averaging the low-resolution 0/1 predictions or doing the same with the 0-1 high-resolution score. To illustrate the sufficiency of the 50 bp low-resolution score when examining larger sets of data in an overview fashion, we sampled 10000 of the promoters used in this study randomly and calculated mean profiles with both the high- and the low-resolution score and compare their similarity in Supplementary Figure S1. The difference between both profiles varies maximally between 0.01 and 0.015. Both resources are easily accessible as bigWig files and listed in the data availability section. Additionally, the code to classify nucleosomal vs. linker DNA can be found there. The NF score described in this section is what is in this work referred to as sequence-intrinsic nucleosome support.

#### DHS score

In this study we analyze the influence of sequence on nucleosome positioning, However, the resulting accessibility in particular cell-lines can be very cell-type specific. Therefore, we created a score to generalize the chromatin accessibility over multiple cell-lines. The DHS (DNase Hypersensitive Sites) score is a measure to estimate the generalized accessibility of any given genome location. For that purpose, we obtained peak-called DNase-seq data for all 403 primary cell lines from ENCODE in human. These we integrated as a simple count of how many primary cell lines show a DNase-seq peak at any given base-pair. To give an example, a score of 403 at a specific bp means that all of the available cell lines show a DNase-seq peak overlapping that position. A score of 5 means that only 5 specific cell lines show accessible chromatin in the form of a DNase-seq peak at the specified bp. In the study, the score is mostly used in the context of indicating TF binding.

## RESULTS

### Sequence-intrinsic nucleosome support characterizes functional genomic regions

We used a random forest classifier to derive a nucleosome support score to estimate the nucleosome support based on the nucleotide sequence and applied it to a range of different genome locations. A broad range of entities can be found in the histone ChIP-seq derived mappings of chromHMM for the cell lines K562 and GM12878 (50). Figure 2 A shows the mean sequence-intrinsic nucleosome support (NF), the nucleosome occupancy (MNase-seq) and the chromatin accessibility (DNase-seq) for the 147 bp in the center of any given chromHMM fragment. In total there are 622257 fragments in K562 and 571339 in GM12878. A list of the individual fragment numbers for each subgroup can be found in Supplementary Table S4. We grouped all chromHMM states into the three categories: “Regulatory” (Promoters/ Enhancers), “Heterochromatin/Repetitive” and “Transcription” (Transcriptional Transition/Elongation, etc. and Insulators).

**Figure 2.**
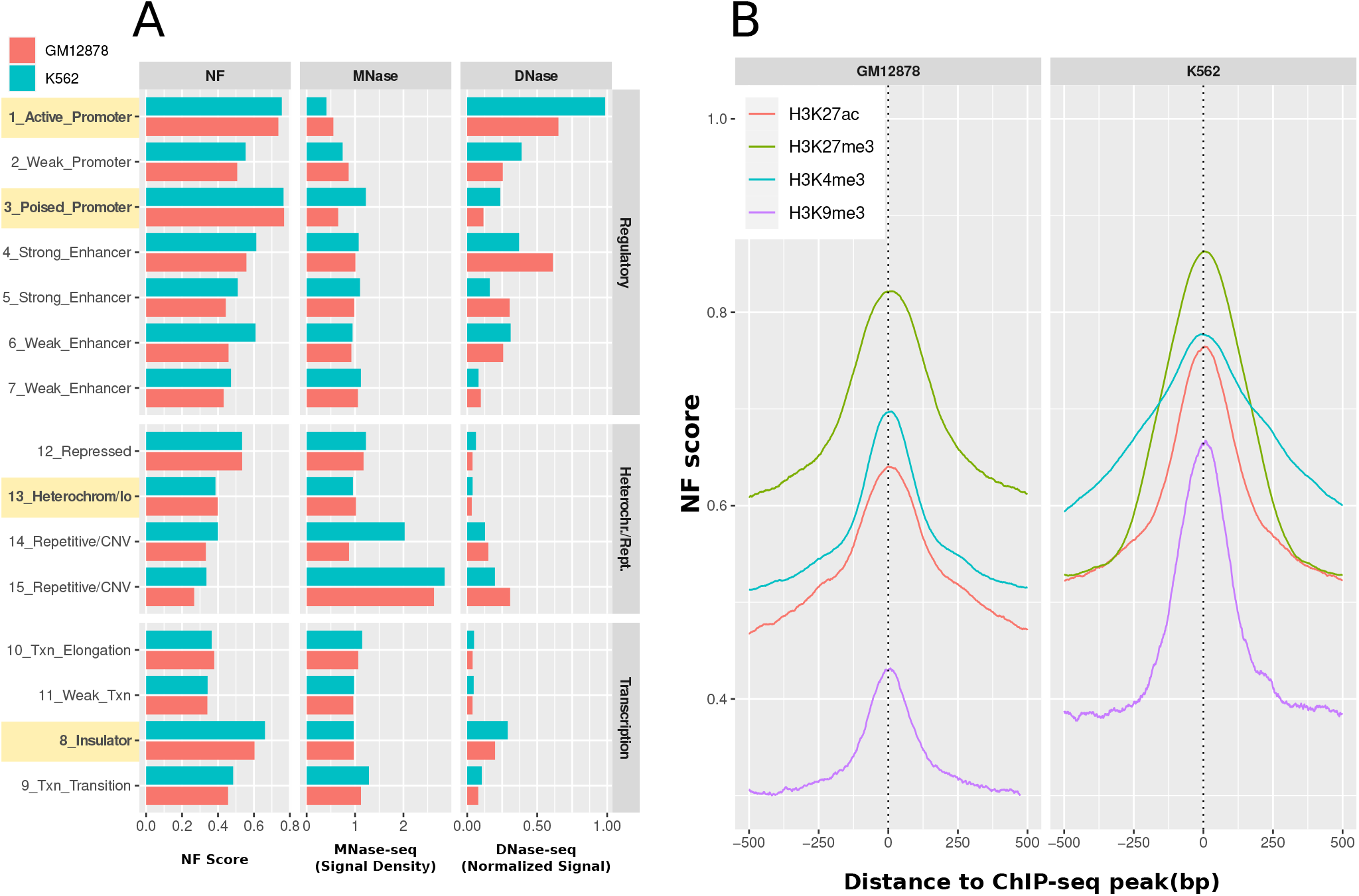
Nucleosome support in different regions of the genome and around histone modification ChIP-seq peaks. **(A)** chromHMM. Nucleosome Support (NF score), nucleosome occupancy (MNase-seq) and chromatin accessibility (DNase-seq) in all states of the chromHMM mappings. The highlighted regions exhibit a high NF score in active regulatory regions (active and poised promoters) as well as in insulators, opposed to low nucleosome support regions in heterochromatin. Active promoters are rather accessible and poised promoters are closed off mostly. Heterochromatin exhibits pronounced inaccessibility, whereas insulators demonstrate a hybrid accessibility characterized by relatively elevated levels of both DNase-seq and MNase-seq signals. **(B)** Nucleosome support of histone modifications. A distinct peak is observed in proximity to each histone modification ChIP-seq peak, providing evidence of the influence of DNA sequence on the positioning of individual nucleosomes. This influence is higher in marks indicating a regulatory function like poised promoters (H3K27me3, green), active promoters (H3K4me3, blue) and more generally accessible chromatin (H3K27ac, red) as compared to repressing histone marks (H3K9me3, purple).

The most intriguing feature is the relation between sequence-intrinsic nucleosome support and actual nucleosome occupancy. Active promoters show the highest DNA-intrinsic nucleosome support (NF: approx. 0.8). However, being crucial places for transcriptional regulation, they are also the most accessible regions (low MNase-seq of 0.4-0.55 and high DNase-seq of 0.65-1). The same nucleosome support can be observed in poised promoters, but with closed chromatin (low DNase-seq, high MNase-seq). This observation can be put in contrast to the highlighted group of heterochromatin fragments. Here, we observe rather low nucleosome support in both cell lines of roughly 0.4. At the same time the locations are very inaccessible with a nucleosome occupancy of 1 and very low DNase-seq signal of 0.03.

The last highlighted group in Figure 2 A are insulators. In this group we can observe a relatively high nucleosome support of 0.6-0.66 and a medium accessibility with relatively high values for both MNase-seq of 1 and DNase-seq signal of 0.2-0.3.

As a side note, it can be observed that enhancers show the most variability in their status. This effect on the NF score is most prominent in the groups of Weak Enhancer (group 6 and 7), in which the NF score varies almost about 0.2 between the minimum in GM12878 and the maximum in K562. These groups are the most ambiguously defined ones, however, based on their histone marks (50). Nevertheless, also in the Strong Enhancer group (group 4) and the Active Promoter group (group 1), there is a comparatively high difference in chromatin accessibility. This could be due to experimental biases but since the direction of the differences is not the same in both instances, it appears that open promoters are generally favored in K562, while the enhancers are more accessible overall in GM12878.

In summary, these observations suggest that regions involved in active regulation of transcription have significantly higher levels of nucleosome support with well-defined sequence context. Whether these important locations are actually occupied by nucleosomes is dependent on whether they are actually used *in vivo* (Active vs. Poised Promoters). In comparison, constitutive heterochromatin has a much lower sequence-intrinsic nucleosome support while being mainly inaccessible.

Histones can carry different kinds of modifications that determine their function and the flexibility of the surrounding chromatin accessibility (14, 15). Since the chromHMM mapping is derived from ChIP-seq data of histone modifications, we can gain more detailed insights into the local nucleosome support relative to the actual position of the histone modification. Therefore, we analyzed 518609 histone ChIP-seq peaks from ENCODE for the GM12878 and K562 cell lines. We chose the histone marks according to their role in relation to the derived chromHMM states. The H3K4me3 peaks are representative for active promoter histone marks (GM12878: 194130, K562: 240828), H3K27ac for enhancers (GM12878: 114738, K562: 102352), H3K27me3 represents poised promoters (GM12878: 100266, K562: 477102) and H3K9me3 is repressive chromatin as a general indication for constitutive heterochromatin (GM12878: 37756, K562: 11778). Whenever there are multiple equivalent experiments for the same target, we calculated the mean over all experiments in a single line.

Figure 2 B shows the average NF score around the set of different histone modification ChIP-seq peaks. It can be seen that, in the specific experimental configurations, the previous findings of a higher NF score in regulatory important regions can be confirmed. For both cell lines, it can be observed that the three activating marks are higher than the heterochromatin mark. Additionally, all histone modifications show a clear peak of nucleosome support around the central ChIP-seq point-source peak. The poised promoter mark H3K27me3 is consistently the most sequence-positioned in both cell lines and can be found at NF score levels of 0.82 and 0.86, respectively. The next highest pattern on both sides is the active promoter mark H3K4me3, which shows a peak at an NF score of 0.67-0.78. The general accessibility mark in promoters and enhancers H3K27ac is next at 0.64-0.76 and the peak of the heterochromatin mark H3K9me3 is located at the lowest NF score level of 0.44-0.66. These results indicate a local sequence preference for nucleosome binding around the central peak position of the histone modifications with a distinctly higher nucleosome support for regulatory elements. Since the modified histones are part of the nucleosome, it can indeed be expected that the NF score is highest around its central binding places but it is remarkable that the sequence-intrinsic influence is highest at putative positions of interaction between nucleosomes and proteins such as TFs.

### Regulatory Nucleosomes: Competition between sequence-intrinsic nucleosomes and transcription factors

#### Promoter regions

The previous analyses suggest that DNA structure plays different roles in nucleosome binding in different biological environments. To enhance our comprehension of these distinctions, we direct our attention towards regulatory loci, specifically encompassing all human promoters. This analytical approach allows us to establish correlations between the unchanging attributes of sequence-intrinsic regulatory potential and its consequential influence on gene regulation. The promoters are defined as 1 kb upstream of all 30142 RefSeq defined human genes (see methods). Alternative TSS locations are counted as individual promoters and strand direction is aligned to 5’-3’ direction.

We applied the high-resolution NF score (see methods) to all human promoters and compare this cell-type unspecific signal with the actual experimental evidence of chromatin accessibility in K562 and GM12878, measured by ENCODE MNase-seq, DNase-seq and H3K4me3 ChIP-seq data as well as a derived DHS score which integrates DNase-seq peaks over 403 primary cell lines (see methods). Additionally, we include PhyloP, a bp resolution evolutionary conservation score.

Figure 3 A shows the position-specific mean distribution for all available genes in relation to the TSS. An extended version of this figure showing 1 kb up- and downstream of the TSS can be found in Supplementary Figure S2. The overall trend of the different data sources is very similar between both cell lines and confirms the general expectations of chromatin accessibility. While the general occupation with nucleosomes declines towards the TSS (MNase-seq from 0.9 to 0.5), the amount of promoter histone modification H3K4me3 increases towards the TSS (from 10 to 25 in K562, from 4 to 7 in GM12878) with a distinct valley about 100 bp upstream of the TSS. This decline in nucleosome occupancy can be explained simultaneously with the gain of chromatin accessibility by putative proteins such as TFs, which is described by a peaked rise of DNase-seq (from 0.25/0.3 to 1.0/1.6) and DHS score (from 10 to 145) in the same area. The NF score increases towards the TSS with a more steady, less peaked slope from approx. 0.5 at −1 kb to 0.85 at the TSS. The evolutionary conservation (PhyloP) stays relatively stable at a level of 0.09 until it increases steeply to 0.45, starting at −200 bp and continuing with a high evolutionary conservation downstream into the gene.

**Figure 3.**
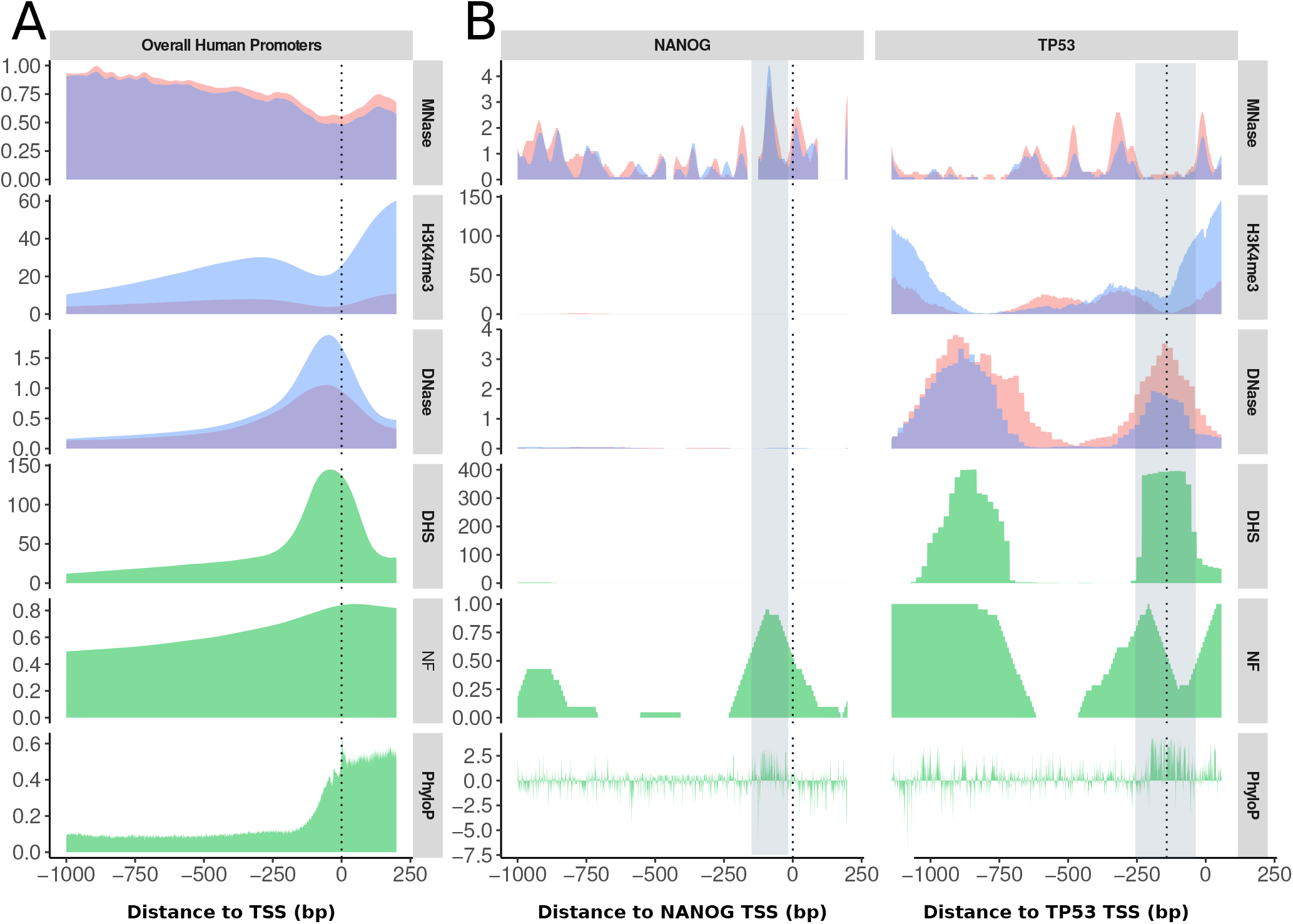
Profiles for experimental chromatin accessibility (MNase-seq, DNase-seq, H3K4me3 ChIP-seq) and static data (green: DHS score, NF score, PhyloP) as (A) the mean of all human RefSeq promoters and (B) the promoters of the individual genes NANOG and TP53 in K562 (blue) and GM12878 (red). The dotted line at position 0 marks the TSS. **(A)** Overall Promoter Regions. The chromatin accessibility increases, while the general nucleosome occupancy decreases. The sequence-intrinsic nucleosome support (NF score) increases towards the TSS along the increasing chromatin accessibility. Furthermore, the H3K4me3 ChIP-seq and MNase-seq signals exhibit an NDR upstream of the TSS and the evolutionary sequence conservation represented as PhyloP score, is generally decreased in the promoter region but increases steeply towards the TSS. **(B)** *NANOG Profiles*. In the context of the two cell lines, NANOG, due to its role as a developmental pioneer factor, does not exert any influence in K562 and GM12878. The marked area denotes an overlap of high sequence-intrinsic nucleosome support as well as a high MNase-seq peak. That means the main site of nucleosome attraction potential is occupied by well-positioned nucleosomes. *TP53 Profiles*. TP53 is an important tumor suppressor in both cell lines. The NF score follows the peaks of high accessibility in DHS score and DNase-seq in location. In the annotated instance, the region characterized by an elevated nucleosome support exhibits a nucleosome-free state *in vivo*, indicating a displaced nucleosome position. Both gene promoters show an increased PhyloP score at the high NF score sites, indicating an evolutionary conservation of the nucleosome supporting sequence. Overall, it should be noted that the NF score correlates positively with the accessible chromatin data (DHS, DNase-seq) in actively used promoters but correlates with nucleosome positioning data (MNase-seq) in closed promoters, demonstrating the original nucleosome positioning potential by the DNA sequence which is overwritten upon promoter activation. *Y-Axes: MNase - density graph of signal enrichment; H3K4me3 - control normalized tag density; DNase - read depth normalized signal; DHS - DHS score, see Methods; NF - NF score (high resolution); PhyloP - PhyloP score)*.

The average profiles of each signal in Figure 3 A are generally coherent, primarily illustrating the concentration of DNA accessibility in proximity to the TSS within the nucleosome depleted region (NDR). However, to understand the local interaction of nucleosomes and TFs, it is crucial to examine the specific positions within the entire promoter region. Figure 3 B shows examples of individual promoters and the relations between nucleosome binding, chromatin accessibility and the potential nucleosome support described by the NF score.

The transcription factor NANOG is a general developmental factor involved in the proliferation and renewal of embryonic stem cells, (62) and thus it does not play a role in the differentiated cell lines. Figure 3 B (left) displays the integrated profiles for the NANOG gene, which is not expected to be actively transcribed in either cell line. Therefore, the first significant observation is the lack of clear signals in both DNase-seq and H3K4me3 ChIP-seq data. Additionally, the general accessibility in primary cell lines, described by the DHS score, is mainly zero overall, too. Albeit, the NF score is high near the TSS (highlighted region) with an NF score of 0.9-0.95. At the same location a maximum of nucleosome occupancy (MNase-seq of 4.4 in K562 and 3.6 in GM12878) can be observed. The evolutionary conservation at this point is much higher with up to 2.9 compared to the near-zero level in the rest of the promoter. This indicates a regulatory nucleosome position that is currently blocked by a well-positioned nucleosome, so that the activation of the NANOG gene requires an active displacement of this nucleosome.

In contrast, the transcription factor TP53 carries substantial importance in both the K562 and GM12878 cell line, as it functions as a critical tumor suppressor (63). Thus, this gene stands as a representative case of a promoter subject to positive regulation. The profiles for this TF can be found in Figure 3 B (right). The accessible regions are separated into two distant peaks on the level of DNase-seq and DHS score, one of which is located around the TSS and the other one at 800 bp upstream. Both accessible regions seem to be important in almost all primary cell lines (DHS score of 401 at the upstream peak and 398 downstream). Both of these peak areas furthermore exhibit an absence of nucleosomes in these areas, evident in an MNase-seq signal of 0-1 at the upstream peak and of 0-0.3 in the downstream peak. Interestingly, the downstream peak is flanked by a high nucleosome occupancy of up to 1.5-2.1 to either side, potentially marking shifted nucleosomes and/or the well-described +1 and −1 nucleosome downstream of the TSS. The sequence support for nucleosome binding in the form of NF score shows a broader plateau of 1 at the upstream portion of the accessible promoter and another maximum directly upstream of the TSS in the NDR with a sharper profile and a short decline before it peaks again closely downstream of the TSS.

Figure 3 A and 3 B together lead us to say that locations in promoters where TFs compete with nucleosomes directly are the places with the highest nucleosome support. Whether these are occupied by nucleosomes is dependent on whether the gene is actually expressed. Thus, we are not of the opinion that all nucleosomes in the promoter are sequence-positioned, as suggested in the mean of all promoters in 3 A. In these overall promoter regions the NF score stays constantly relatively high but in the individual promoters in 3 B, it depends on the co-localization with competing TFs that mediate chromatin accessibility at local, defined places.

#### Transcription Factors

The regulation of gene expression is highly dependent on the binding of TFs. TFs exhibit a range of strategies in their competition with nucleosomes, displaying remarkable adaptability. Nevertheless, a common feature among them is that a region occupied by TFs should typically be devoid of nucleosomes. To give an overview of the nucleosome support for the diverse array of TFs, we calculated the mean NF score for all available ENCODE TFs at the peak positions of optimal IDR threshold peaks. The number of available files with at least two biological replicates is 151 for GM12878 and 391 for K562. Figure 4 A shows a histogram of NF scores at each according individual peak point position as defined in the Methods section. It is notable that the overall mean nucleosome support of TFs is in general rather high with approx. 0.71 and very consistent between the two cell lines with a difference within less than 1%. The difference in appearance between these two groups lies solely in the imbalance in the number of available experiments for each cell line. Furthermore, we calculated the mean NF score around the TF ChIP-seq peak location summarizing all of the TFs into one profile which is depicted in Figure 4 B. The peak of the NF score coincides precisely with the center of the experimentally validated ChIP-seq peaks. As expected, the NF score at this position matches precisely with the median NF score at the peak position depicted in Figure 4 A’s histogram, measuring 0.71 at the peak’s maximum position. The figure illustrates that, in the vicinity of TF binding events, the MNase-seq signal experiences a consistent reduction (GM12878: signal decrease from 0.91 to 0.85; K562: signal decrease from 1.01 to 0.89). In contrast, the NF score reaches its peak precisely at these locations, suggesting the presence of sequence-intrinsic nucleosome positioning, particularly in regulatory regions that compete with initially well-positioned nucleosomes. The NF score lies in between what is to be expected for enhancer and promoter regions considering Figure 2. Due to the abundance of ChIP-seq peaks that are expected to be located in enhancer regions, the profiles might be biased towards enhancer TFs. In order to show whether promoters and enhancers behave differently, we added the information of H3K4me3 and H3K27ac signal for the TF ChIP-seq peaks in Supplementary Figure S3. In this context, it is apparent that both signals exhibit a consistent depletion pattern in close proximity to the binding locations, demonstrating a strong similarity in their behavior.

**Figure 4.**
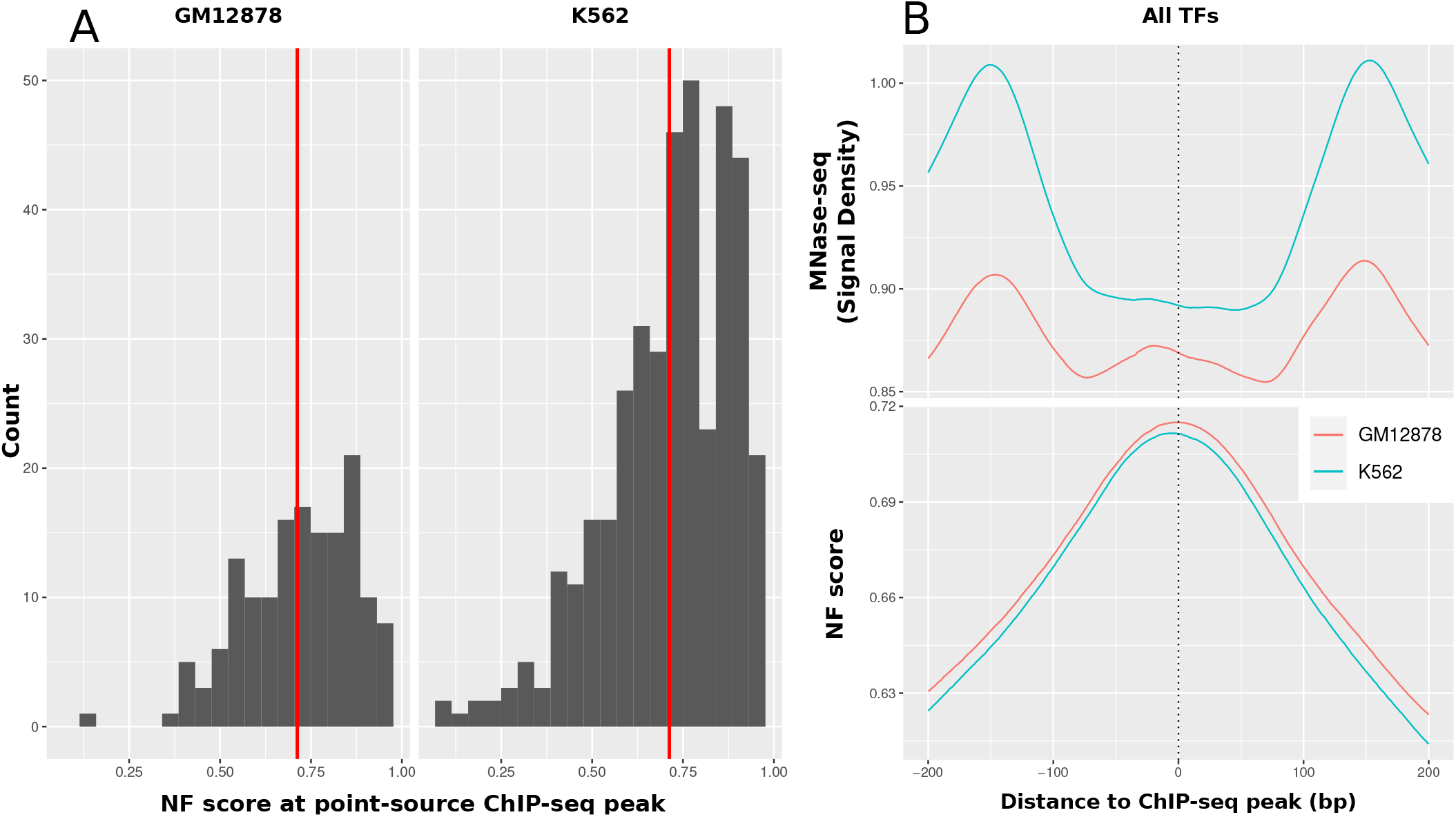
Overall sequence-intrinsic nucleosome support around human TF binding locations. **(A)** Sequence support at TF binding locations. Histogram of the NF score at the point-source peak position of all available ChIP-seq peaks on ENCODE for K562 and GM2878. The red line marks the mean of NF score at peaks (GM12878: 0.7150, K562: 0.7114). **(B)** Average sequence-intrinsic nucleosome support (NF score) around the point-source peak of all TFs in comparison to nucleosome occupation (MNase-seq) in K562 and GM12878. It can clearly be seen that the sequence-based positioning signal is highest around the ChIP-seq peak location (0.71) while this is the position with the lowest nucleosome occupancy *in vivo*. This is evidence for the high importance of sequence to position nucleosomes based on sequence at places where competition with TFs is highest.

Since ChIP-seq only covers the final state of the displacement of the nucleosomes, we illustrate the relation of originally closed off chromatin and the sequence-based nucleosome positioning with a pioneer-factor that serves the purpose of displacing the nucleosome and making the chromatin accessible. The pioneer transcription factor GATA3 has been described to bind well positioned nucleosomes (64). To check whether this positioning can be seen on DNA level as well, we analyzed 43504 GATA3 binding sites with our global NF score and compared them to published nucleosome occupancy (MNase-seq) in MDA-MB-23 cells (47). Figure 5 A shows the profiles of +-500 bp around GATA3 binding sites. It is evident that the area is supportive for nucleosome binding based on sequence as shown by an NF score that shows an increased peak of approx. 0.15 around the binding sites from 0.35 to 0.5. In the case of cells without GATA3 expression (control), there is a shift of increased nucleosome occupancy (0.63-0.7) around the sites, showing increased nucleosome occupancy on top of the closed binding sites. However, upon GATA3 expression, the peak observed in the data undergoes a split into a double peak at 0.675, corresponding to both sites, while exhibiting a distinct valley at 0.62 directly at the transcription factor binding sites (TFBSs). This behavior suggests the presence of a robust nucleosome that was initially positioned based on the DNA sequence but subsequently displaced due to the influence of pioneer transcription factors.

**Figure 5.**
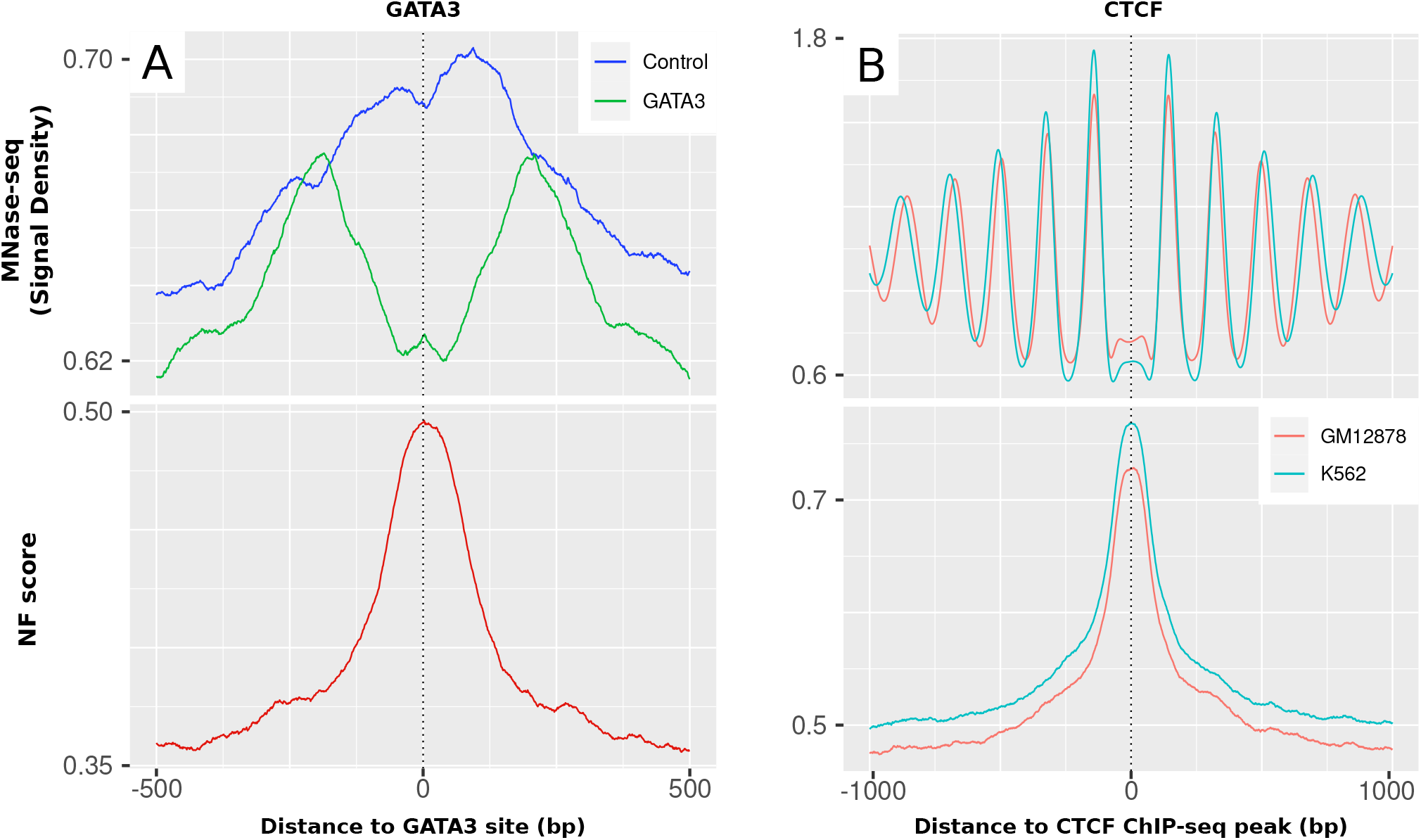
Nucleosome Formation score (NF) and nucleosome occupancy (MNase-seq) surrounding the transcription factors GATA3 and CTCF. **(A)** GATA3. GATA3 binding sites of MDA-MB-23 cells with GATA3 expression (green: GATA3 expression) and without (blue: Control, no expression of GATA3). The nucleosome occupancy is higher towards the binding sites in the cells without GATA3 expression. This peak is depleted in the GATA3 expression cells and a double peak of nucleosome occupancy is left around the sites with a distance of approx. 200 bp. The nucleosome support displays a pronounced peak precisely overlapping the TFBSs, indicating the presence of an intrinsically well-positioned nucleosome that is actively displaced by GATA3. **(B)** CTCF. CTCF ChIP-seq peaks from ENCODE in GM12878 and K562. The nucleosome occupancy shows a very distinct periodicity around CTCF peaks. This periodicity of nucleosome positions is not apparent on the level of sequence-intrinsic nucleosome support, which exhibits a single peak directly around the CTCF peak.

#### Insulators

Another category of regions with a rather high mean NF score of 0.66 in Figure 2 A were insulator regions. Insulators are DNA regulatory elements that play a crucial role in organizing and maintaining the spatial organization of the genome. The protein CTCF is a key factor in mediating insulator function by facilitating chromatin looping and regulating gene expression (65). The nucleosome occupancy and sequence-intrinsic nucleosome support around 43247 ChIP-seq peaks for CTCF in K562 and 43631 peaks in GM12878 are compared in detail in Figure 5 B. The peaks are agnostic of strand orientation, since they are not mapped to a particular gene.

The nucleosome occupancy shows a very stable, phased array of nucleosomes around the CTCF peaks with an approx. 200 bp periodicity and an ongoing MNase-seq signal decrease per peak from 1.8 to 1.2 with increasing distance to the TSS. While the nucleosome occupancy is lowest directly at the CTCF peaks, the sequence-intrinsic nucleosome support is rather high specifically in these regions with an NF score of about 0.77 at the peak. The observed periodicity in nucleosome occupancy is completely absent on the sequence level, indicating a strong influence on nucleosome positioning at CTCF sites but active energy-driven chromatin remodeling once the site was cleared and the factor is bound to the DNA.

### Sequence-intrinsic nucleosome support in transcriptional regulation

In the previous part we focused mainly on the role of DNA for nucleosome positioning in promoters, in proximity to the TSS and in competition to TFs. The following sections will explore other functional elements of the genome along the transcriptional axis in a similar way.

#### Active chromatin remodeling upon transcription initiation

It is known that there is a multitude of influences on the dynamic positioning of nucleosomes. One such phenomenon is the phasing of ordered nucleosome arrays downstream of the TSS upon transcription initiation (66). Figure 6 A shows the comparison of nucleosome occupancy (MNase-seq) and the DNA-intrinsic nucleosome support (NF) around the TSS of all human RefSeq genes. The proposed periodic downstream array of nucleosome positions is apparent, marked by arrows, stretching with a periodicity of 200 bp at MNase-seq levels between 0.75 and 0.9 (in GM12878) or 0.65 and 0.85 (in K562), respectively. At the same time, although the NF score is generally still relatively high close to the TSS and the +1 nucleosome with a value of 0.8-0.85, the periodicity is not mirrored on the sequence level and the general sequence-intrinsic nucleosome support decreases further downstream. Therefore, it is reasonable to conclude that the dynamic re-localization of nucleosomes due to ATP-dependent remodeling can overwrite the sequence-mediated nucleosome guidance.

**Figure 6.**
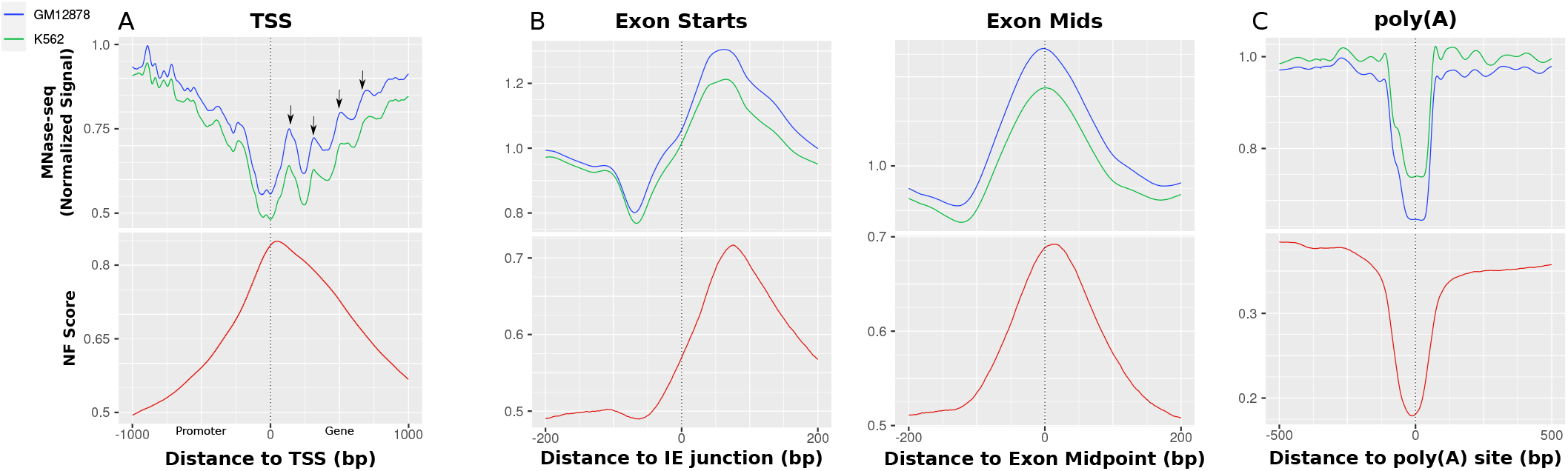
MNase-seq and NF score for different elements from transcription initiation to transcription termination. **(A)** Promoters. The profiles are consistently aligned to ensure that the promoter located at the 5’ end is consistently positioned on the left side. The central vertical line represents the TSS location. The nucleosome occupancy declines around the TSS and shows an apparent periodicity in both directions. The NF score rises in anti-correlation towards the TSS and does not indicate the proposed periodicity. This implies there is a support for nucleosome binding for the NDR which is accessed by TFs but no sequence support that places the periodically following nucleosomes in the downstream array. **(B)** Exons. The left image is centered specifically on the Intron/Exon (IE) junction, while the image on the right aligns the exons precisely over their midpoints. The NF score shows a high sequence-intrinsic nucleosome support and an experimentally high nucleosome occupancy directly downstream of the exon starts, which can be observed even more stably over exon centers. This correlation strongly suggests the functional significance of the observed sequence-defined nucleosome. **(C)** Poly(A). There is a decline of both sequence-intrinsic nucleosome support (NF score) and nucleosome occupancy (MNase-seq) around the transcription termination sites. This behaviour indicates nucleosome-repelling positions, encoded in DNA sequence.

#### Sequence-intrinsic nucleosome occupation potential of exon regions

While it was just shown that there is a sequence-independent creation of nucleosome arrays downstream of the TSS, that does not entirely describe the sequence-intrinsic local nucleosome support for other regions along transcriptional regulation. It is described in literature that exons are targets of a particularly high nucleosome occupancy which is related to transcription speed of RNAPII and the recognition of alternative splicing (11). To test whether this principle is also evident on the nucleotide sequence level, we analyzed the exons of all human genes by comparing the MNase-seq signal for K562 and GM12878 and the NF score again. Figure 6 B shows both average signals centered around 267864 exons with strand orientation taken into account. The figure on the left employs an alignment centered around the Intron/Exon (IE) junction, specifically focusing on the exon starting points. Conversely, the right-hand figure emphasizes the alignment based on the central exon midpoints. Generally, the figures indicate that there is a nucleosome positioned over the center of the exon. The midpoint-centered figure shows that the nucleosome occupancy and the NF score show a distinct peak over the exon regions. The extent of the rise of the NF score towards the exon center is as high as approx. 20% from slightly above 50% to almost 70% and is distributed mainly between 100 bp upstream and 150 downstream of the exon midpoints. The nucleosome occupancy mirrors this pattern of a central peak of about 0.3-0.4 from 0.9 up to 1.25 in both cell lines around the exon. In addition, a notable observation indicates a relative depletion of MNase-seq signal directly upstream of the peak, which is more clear when aligning the exons over the IE junction. In this view, it is also apparent that the main nucleosome is positioned directly downstream of the IE junction. That the middle-centered nucleosome can be understood as equivalent to the nucleosome directly downstream of the exon start can be explained with the median of the RefSeq exons length being 129, thus being not large enough to fit multiple nucleosomes. The general tendency of a single sequence-positioned nucleosome in the center of an exon holds true when separating the exons by length. For a more detailed analysis of these patterns, Supplementary Figure S4 and Supplementary Figure S5 provide an enhanced representation by separating the exons based on their length, again around the midpoints and around the IE junction, respectively.

#### Sequence-dependent nucleosome occupancy prevention around poly(A) sites in transcription termination

Another important part of gene regulation and splicing is the nucleosome occupancy at poly(A) sites as the place for transcription termination. These were shown to be explicitly free of nucleosomes, thus making them accessible for proteins marking the sites and interacting with RNAPII in transcriptional termination (13). We obtained the ENCODE MNase-seq signal for K562 and GM12878, as well as the NF score, for a list of 570740 poly(A) sites in the human genome (52). These are classified by the original authors into clusters of biological origin. All sites are oriented by strand in 5’ - 3’ direction. Figure 6 C shows the nucleosome occupancy and NF score for all poly(A) sites in the mapping together. There appears to be a direct positive relation between the sequence-intrinsic nucleosome support and the nucleosome occupancy. Both nucleosome occupancy and sequence-intrinsic nucleosome support exhibit a decrease around the sites in the central 250 bp. There is a 20% drop in nucleosome support relative to the already low level of 35%. It can be noted that the sequence support for nucleosome binding seems to be slightly higher in the upstream direction. The same overall decrease is apparent on nucleosome occupancy in both cell lines with a drop of 0.25-0.3 in MNase-seq signal. A proposed increase in nucleosome occupancy downstream of the poly(A) site *in vivo* relative to the upstream direction (13) can not be observed overall. Nevertheless, outcomes for specific subgroups of poly(A) sites can differ. Supplementary Figure S6 gives an overview over all poly(A) sites separated by clusters defined in the PolyASite 2.0 atlas. There, it can be observed that the general repelling trend can be observed for all subclusters, except for sites that coincide with exon regions, which exhibit a similar pattern as shown above for exons, i.e., a sequence-positioned nucleosome over the exon. The aforementioned difference of a higher nucleosome occupancy downstream of the site can be shown there for the DS cluster, which denotes the TTS 1 kb downstream of a terminal exon. In contrast, the higher NF score in the mean profile upstream of the sites can most probably be explained by the group of sites located in the AU group, in which the signal is higher due to its proximity to the TSS which has been demonstrated to have a rather high NF score in Figure 6 A.

## DISCUSSION

In this study, we analyzed the role of DNA sequence for the positioning of nucleosomes. For that purpose, we developed a random forest classifier to distinguish nucleosomal and linker DNA and used it to derive a score to estimate the sequence-intrinsic nucleosome support of different functional elements in the human genome. The degree to which nucleosome positioning relies on DNA sequence has been the subject of extensive debate (16). These questions cannot be answered by analyzing either chromatin accessibility experiments or DNA sequence individually. On the one hand, the experimental verification of chromatin accessibility for any given cell-type will only describe the result of the multitude of counteracting influences on nucleosome positions. On the other hand, DNA can explain the importance of nucleotide-sequence on their potentially favoured binding locations, without reliably predicting the actual nucleosome occupancy in any given cell-type *in vivo*. Therefore, we compared the results of our sequence-intrinsic nucleosome support in the form of the NF score with experimental data for chromatin accessibility and histone modifications and described the possible combinations of *in vivo* nucleosome positioning with its underlying DNA sequence support. We could demonstrate that DNA-intrinsic nucleosome support is not directly equivalent to the resulting nucleosome occupancy in a specific cell-line *in vivo*. Rather, there is a positive influence on nucleosome binding in the DNA sequence of genomic regions which are involved in actively regulating transcriptional processes, such as promoters, enhancers, exons and in the center of insulators. Simultaneously, certain locations exhibit lower nucleosome support, which can serve functional purposes, such as well-positioned nucleosome occupancy arrays downstream of the transcription start site (TSS) and in the vicinity of insulators. Conversely, there are instances where lower nucleosome support indicates non-functionality, as observed in constitutive heterochromatin.

The increased reliance on DNA sequence in gene regulatory regions could be shown in this study from a variety of biological angles. The apparent general trend in promoter regions could be narrowed down to be due to the direct places of competition with TFs. The nucleosome support by sequence could be attributed to evolutionary pressures favoring the development of DNA sequences that facilitate optimal nucleosome binding, effectively regulating access to these regions. This effect has been described to be particularly important for organisms of higher complexity (61). Especially these highly sequence-intrinsic and evolutionary conserved positions can make a large difference in the regulation of gene activity and thus, the sequence can help to increase the resistance to accidentally triggering gene expression. The counteracting process to this state is an active eviction of the blocking nucleosome by a combination of (pioneer-)TFs, histone modification enzymes and active chromatin remodeling to gain accessibility for a particular position (67). Due to the inclusion of the relevant TFBSs within the confined nucleosome positions, it can be inferred that there is a presumed co-evolution of proficient nucleosome organization and the existence of high-quality binding sites for activating TFs. Therefore, we interpret the crucial function of DNA in these “regulatory” nucleosomes as a mechanism to regulate the competition between nucleosomes and TFs for spatial availability. This competition affects the energy required to activate biological processes, allowing for dynamic control and modulation. Further evidence for the systematic co-localization of regulatory proteins (high DHS score) with highly nucleosome-attracting regions (high NF score) can be observed in Supplementary Figure S7. That the accessibility of these sequence-defined nucleosomes plays a significant role in regulating gene expression is further supported by Supplementary Figure S8, in which gene expression values are compared between groups of genes based on a correlation or anti-correlation of NF score and chromatin accessibility data. The role of DNA sequence for different kinds of TFs under this assumption has been partially analyzed for different TF families (68, 69). Nevertheless, it would be worthwhile to research the differences in sequence-intrinsic nucleosome support in regard of pioneer functionality. The co-evolution to occupy highly important places would suggest a high nucleosome support at pioneer TFBSs. However, this presents a notable challenge as the binding sites of pioneer transcription factors are often located in close proximity to the subsequent TFs that bind in a consequent step and that are not directly in competition with the nucleosome. Furthermore, it is well-established that pioneer TFs can employ various modes of nucleosome eviction, which may not occur directly at the target site of chromatin accessibility (7, 70). Finally, while there is a difference in the set of TFs that are responsible for regulating the chromatin accessibility of promoters and enhancers (71), we could show that for the general relationship of nucleosome positioning in the competition with TF binding, both groups show a similar behaviour.

In contrast to these nucleosome positions determined by DNA sequence, constitutive heterochromatin, although commonly occupied by nucleosomes, does not demonstrate a comparable reliance on DNA sequence for precise nucleosome positioning. In analogy to the previous explanation of DNA evolution towards a targeted occlusion of TFBSs to prevent random gene activation, we propose a lower relevance of sequence for heterochromatin. The higher-order packing of dense chromatin is a more unspecific process in which the accessibility of small individual positions on the genome is not expected to have such severe consequences as the revealing of TFBSs in cis-regulatory elements.

Apart from the evolution of nucleosome positioning advantage at highly competitive sites, we found other instances of locations that appear to have a high sequence-intrinsic nucleosome support. In particular, exons show a considerable nucleosome support relative to their surroundings. It is well described that the occupation of exon regions with nucleosomes is important in the pausing of RNAPII to ensure the correct transcription of a gene as well as the marking and spatial relocation of exons for the splicing machinery (11, 72). Certain studies propose that the sequence specificity for nucleosomes at exons is primarily encoded at the exon’s outset, specifically in the form of a specific dinucleotide distribution at the Intron/Exon junction (73). However, the positioning of the nucleosome based on the sequence is found in this study directly downstream of the exon start. This finding is in line with the theory that the nucleosome in the exon stalls the RNAPII just upstream of the IE junction (74). The principle of using the DNA to encode a favourable structure to guide nucleosomes to exon regions might again be an efficient means of increasing the intrinsic nucleosome support. In contrast to strengthening the nucleosome in competition with TFs, the nucleosome at exons might simply be guided to its place of effect without the need for further displacement by other processes. The difference to the non-sequence-dependent nucleosome positioning at the TSS could be grounded in the reduced energy needed to stabilize nucleosomes permanently at exons compared to the potential for dynamic shifting of nucleosomes around TSS upon gene activation. This provides another striking example of putative co-evolution that is directed to attract nucleosomes on the sequence which underlies a drive to reliably encode proteins at the same time.

The utilization of sequence to influence nucleosome positioning is evident not only in its ability to positively guide nucleosomes to specific locations. We showed the opposite effect for poly(A) sites, marking the end of transcription. The substantial contrast in sequence-intrinsic nucleosome support between poly(A) sites and the surrounding DNA clearly indicates the strategic prevention from nucleosome binding at these specific locations. The reason for this feature could be the attempt to keep chromatin at the TTS constantly accessible to easily allow the binding of proteins that form complexes together with the RNAPII to terminate transcription, cleave the mRNA and do the polyadenylation in the consecutive step. The nucleosome rejection fits well to the general observation of low nucleosome occupancy over poly(A) sites. However, there are hints about the role of increased nucleosome occupancy around one end of the sites, regulating alternative polyadenylation and depending on status of active transcription (13). This could be the reason for a difference of nucleosome support towards both directions surrounding poly(A) sites in terminal exons. Here, we observed a slightly lower nucleosome support downstream of poly(A) sites for terminal exons but not as a general trend over all groups. Also we could show an increased nucleosome support upstream of the TSS for the group of sites downstream of the TSS, in which the increase is easily explainable by the overlap with the generally sequence-defined promoter region. It cannot definitively be determined whether the otherwise relatively constant nucleosome occupancy up-and downstream arises from technical aspects in the normalization of ENCODE MNase-seq data or if it is attributed to variations in the splicing patterns of different clusters of poly(A) sites, particularly when situated in regions other than terminal exons, or whether all of these sites possess transcription termination functionality at all.

Opposed to the steering of functional nucleosomes by nucleotide sequence, we identified processes which exhibit strong nucleosome positioning that does not manifest on DNA sequence level. Notably, downstream of the TSS, a well-established pattern of regularly spaced nucleosome array formation is observed. At these locations the nucleosome support constantly decreases towards intronic locations without mirroring the periodicity of the nucleosome occupancy. This process is known to be mediated by active chromatin remodeling complexes which are organizing nucleosomes into an evenly spaced array by using energy from ATP (10). This process serves as an example for an important transcriptional mechanism which is not directly supported by local DNA structure. Regarding the nucleosome arrays adjacent to the TSS, it can be speculated that the +1 and −1 nucleosomes are influenced by sequence-specific influences. This assertion is supported by the relatively high NF score, aligning with studies showing the special significance of these particular nucleosomes (8, 75). These studies suggest a discernible dinucleotide enrichment in proximity to these nucleosome locations, implying a mechanism of “statistical positioning” for the co-occurring nucleosome arrays. However, the proximity of these nucleosomes to the main areas of TF-nucleosome competition in the promoter could easily lead to misinterpretation of their reliance on DNA sequence alone.

In conclusion, it is worth highlighting that these processes of sequence-intrinsic and sequence-independent nucleosome positioning may not necessarily be mutually exclusive. This phenomenon is particularly pronounced at insulator sites, where the transcription factor CTCF plays a crucial role in delineating TAD boundaries. Here, it could be observed that on top of the actual binding location there is a very strong sequence-defined support for the nucleosome, suggesting the previously described mechanism of competition and co-evolution of TF-nucleosome interaction. Yet, as a result in the surroundings of CTCF bound locations, there is a clear well-positioned nucleosome array which is the result of active chromatin remodeling without the help of sequence support. While the binding of CTCF underlies the aforementioned competition with positioned nucleosomes, the spacing and symmetry of the surrounding array is rather linked to active chromatin remodelers, instead (5). This is just one example of the very close, local interplay of sequence support and energy-dependent remodeling of chromatin to decide the final *in vivo* positions of nucleosomes in transcriptional regulation. It is impossible to favour any of these principles in their importance for the collaborative regulation of transcription, from the formation of TAD boundaries, over the regulation of gene activity through the accessibility of chromatin for TFs or support mechanisms of the tasks of RNAPII, to the clean termination of transcription, thus marking the last step in which the DNA is involved directly.

## DATA AVAILABILITY

The code to execute predictions can be found at https://gitlab.gwdg.de/MedBioinf/generegulation/nfclassifier. The NF scores in bigWig file format can be downloaded at https://bioinformatics.umg.eu/resources/nfscore.

## ACKNOWLEDGEMENTS

We thank Argyris Papantonis and Alexander Ecker for fruitful discussions. MS is a member of the International Max Planck Research School for Genome Science (IMPRS-GS), part of the Göttingen Graduate Center for Neurosciences, Biophysics, and Molecular Biosciences (GGNB). NP is a member of Molecular Medicine PhD program, part of the Göttingen Graduate Center for Neurosciences, Biophysics, and Molecular Biosciences (GGNB). TB and MH are members of the Göttingen Campus Institute Data Science (CIDAS). Also, we thank the ENCODE Consortium and the ENCODE production laboratories generating the particular data sets.

