## Supplementary Material for "The importance of DNA sequence for nucleosome positioning in transcriptional regulation"

**Supplementary Table S1.** ENCODE IDs for individually mentioned datasets used in this study.

| | Experiment | Cell | ENCID |
|---|---|---|---|
| 1 | MNase-seq | K562 | ENCFF000VNN |
| 2 | MNase-seq | GM12878 | ENCFF000VME |
| 3 | DNase-seq | K562 | ENCFF352SET |
| 4 | DNase-seq | GM12878 | ENCFF901GZH |
| 5 | H3K4me3 (location) | K562 | ENCFF378OQB,ENCFF883POK,ENCFF127XXD,ENCFF737AMS |
| 6 | H3K4me3 (location) | GM12878 | ENCFF795URC,ENCFF354MGT |
| 7 | H3K27me3 (location) | K562 | ENCFF322IFF,ENCFF126QYP |
| 8 | H3K27me3 (location) | GM12878 | ENCFF851UKZ,ENCFF247VUO |
| 9 | H3K9me3 (location) | K562 | ENCFF894VEM |
| 10 | H3K9me3 (location) | GM12878 | ENCFF682WIQ |
| 11 | H3K27ac (location) | K562 | ENCFF044JNJ |
| 12 | H3K27ac (location) | GM12878 | ENCFF816AHV |
| 13 | H3K4me3 (signal) | K562 | ENCFF285EUA |
| 14 | H3K4me3 (signal) | GM12878 | ENCFF154XCY |
| 15 | H3K27ac (signal) | K562 | ENCFF488FYZ |
| 16 | H3K27ac (signal) | GM12878 | ENCFF440GZA |
| 17 | CTCF (location) | K562 | ENCFF002DDJ |
| 18 | CTCF (location) | GM12878 | ENCFF710VEH |

**Supplementary Table S2.** ENCODE IDs of all ChIP-seq experiments used for the TF analysis in GM12878.

**GM12878:** ENCFF814CZI, ENCFF134XDY, ENCFF711TMS, ENCFF229YSW, ENCFF123WEZ, ENCFF809PXE, ENCFF344OJI, ENCFF456FQB, ENCFF935HIG, ENCFF881VZH, ENCFF711OSR, ENCFF103OOV, ENCFF381SBC, ENCFF456YII, ENCFF349MPF, ENCFF199WGD, ENCFF817QHW, ENCFF557HJB, ENCFF909FRA, ENCFF917CXD, ENCFF788LSH, ENCFF339FFC, ENCFF208TRU, ENCFF895YBG, ENCFF827ZLB, ENCFF116EXQ, ENCFF980KSV, ENCFF760IFZ, ENCFF692GBM, ENCFF609KOY, ENCFF160WLU, ENCFF079ZWK, ENCFF433PFN, ENCFF020GMZ, ENCFF073XZC, ENCFF273ULT, ENCFF461FGJ, ENCFF630NFM, ENCFF653MIQ, ENCFF717IXP, ENCFF467NRS, ENCFF632MBO, ENCFF478DRD, ENCFF805GKD, ENCFF372DRC, ENCFF303TVZ, ENCFF048JKT, ENCFF997NGJ, ENCFF868VSY, ENCFF420YHY, ENCFF195PBG, ENCFF931XAL, ENCFF967ACD, ENCFF066XDW, ENCFF445UJF, ENCFF733UTZ, ENCFF704ZYN, ENCFF556PIY, ENCFF721SDD, ENCFF394DLH, ENCFF405NFV, ENCFF712GLE, ENCFF277VDY, ENCFF261ZWR, ENCFF242VUO, ENCFF899JBI, ENCFF490VYC, ENCFF142EVF, ENCFF678JII, ENCFF248QFF, ENCFF374HJR, ENCFF339KUO, ENCFF391DVL, ENCFF082JDH, ENCFF556JBS, ENCFF014RBU, ENCFF305ICK, ENCFF534CKB, ENCFF298UPI, ENCFF745TNJ, ENCFF141SAU, ENCFF544NRC, ENCFF199FYD, ENCFF628QJU, ENCFF721FTG, ENCFF778DJD, ENCFF394CWJ, ENCFF853SOB, ENCFF593FBF, ENCFF341HWZ, ENCFF976ZVG, ENCFF409QJU, ENCFF273BVT, ENCFF241GZK, ENCFF735XQT, ENCFF402FYF, ENCFF328QLX, ENCFF468MFY, ENCFF810CEL, ENCFF686FLD, ENCFF957YRU, ENCFF677KJB, ENCFF332PGQ, ENCFF310XJC, ENCFF625SHY, ENCFF083KVY, ENCFF898MUV, ENCFF753RGL, ENCFF030NYG, ENCFF761MGJ, ENCFF643OWO, ENCFF492YMU, ENCFF413VUC, ENCFF516AJC, ENCFF304WER, ENCFF606WUV, ENCFF317VBQ, ENCFF563QBC, ENCFF357WYN, ENCFF127GYQ, ENCFF689AXJ, ENCFF371OGO, ENCFF414JLN, ENCFF055NJR, ENCFF455YHJ, ENCFF063WXY, ENCFF302XID, ENCFF015YHT, ENCFF017LZE, ENCFF455HXN, ENCFF125SQT, ENCFF710VEH, ENCFF166FRO, ENCFF499ZPP, ENCFF107UHX, ENCFF996RRT, ENCFF809OOE, ENCFF911BYP, ENCFF966RPL, ENCFF100ERQ, ENCFF939TZS, ENCFF513RIQ, ENCFF449CJP, ENCFF120VUT, ENCFF833FTF, ENCFF807AKG, ENCFF566YUW, ENCFF222XPQ, ENCFF462GHD, ENCFF002DCM, ENCFF473RXY

**Supplementary Table S3.** ENCODE IDs of all ChIP-seq experiments used for the TF analysis in K562.

**K562:** ENCFF583DZD, ENCFF196DHR, ENCFF213EPU, ENCFF602OKQ, ENCFF861YKK, ENCFF388AJH, ENCFF828JWK, ENCFF232KAH, ENCFF941RVL, ENCFF484DKT, ENCFF646VQW, ENCFF597DIY, ENCFF246HIU, ENCFF859NPS, ENCFF802XZN, ENCFF782GWS, ENCFF165BCW, ENCFF228PSB, ENCFF442ZUF, ENCFF421VCV, ENCFF556YCY, ENCFF671BFH, ENCFF408SNO, ENCFF906PGD, ENCFF677OWM, ENCFF284PKK, ENCFF932UEC, ENCFF684MLP, ENCFF788EBS, ENCFF809YFY, ENCFF226CZE, ENCFF643TEZ, ENCFF495CSO, ENCFF784FIE, ENCFF682WUJ, ENCFF757ODD, ENCFF798URE, ENCFF289BUU, ENCFF292JRY, ENCFF698MLW, ENCFF265LKN, ENCFF996ZGL, ENCFF332DRG, ENCFF262BVP, ENCFF877ANE, ENCFF679ITJ, ENCFF526HLP, ENCFF728NEV, ENCFF583GKE, ENCFF589EVD, ENCFF583CIY, ENCFF859FVX, ENCFF901AHW, ENCFF310AGG, ENCFF272ULO, ENCFF117EWO, ENCFF396DNK, ENCFF493TZM, ENCFF549GMO, ENCFF973OME, ENCFF772QPO, ENCFF447QUG, ENCFF257RYT, ENCFF993GXU, ENCFF209JJD, ENCFF642BNC, ENCFF134CUM, ENCFF594LCH, ENCFF769GYG, ENCFF836LBG, ENCFF473GCH, ENCFF253FZN, ENCFF629BFI, ENCFF772HOY, ENCFF895QLA, ENCFF398TWC, ENCFF558VPP, ENCFF034CQR, ENCFF269EMM, ENCFF059ONJ, ENCFF048OBR, ENCFF803LEC, ENCFF835KAT, ENCFF076YZO, ENCFF308IXJ, ENCFF004WYV, ENCFF352HGW, ENCFF106DAY, ENCFF370ENX, ENCFF122QSN, ENCFF359QCN, ENCFF708BTH, ENCFF033KXY, ENCFF574LAO, ENCFF605KFA, ENCFF970LCB, ENCFF958KNK, ENCFF561IZB, ENCFF799HIG, ENCFF670ZCR, ENCFF278ONE, ENCFF452SVO, ENCFF891MVM, ENCFF696KPD, ENCFF844HBQ, ENCFF922TDM, ENCFF626MVT, ENCFF225MPC, ENCFF834WGJ, ENCFF545RHH, ENCFF474NLG, ENCFF886VSU, ENCFF874QUM, ENCFF493ABN, ENCFF809MSQ, ENCFF163VUK, ENCFF521CRG, ENCFF659WGE, ENCFF273EYJ, ENCFF033EBX, ENCFF806BDC, ENCFF462CPT, ENCFF730URE, ENCFF905VXX, ENCFF042RAK, ENCFF722XRW, ENCFF241TBP, ENCFF567GSX, ENCFF076MSV, ENCFF881QBT, ENCFF492GXZ, ENCFF426DUB, ENCFF662MVX, ENCFF564WEB, ENCFF314ULQ, ENCFF248VHN, ENCFF972LPT, ENCFF484CKD, ENCFF581ZZT, ENCFF581YKY, ENCFF522JRK, ENCFF422BXE, ENCFF968JVX, ENCFF380FJL, ENCFF011KRN, ENCFF870PWV, ENCFF315YAF, ENCFF344QKL, ENCFF962FQM, ENCFF517KRT, ENCFF803COK, ENCFF710LRU, ENCFF632NQI, ENCFF059WVE, ENCFF246VJH, ENCFF671DOL, ENCFF557DSM, ENCFF706ISJ, ENCFF710IEF, ENCFF152VMJ, ENCFF701NVT, ENCFF392MUM, ENCFF417LEJ, ENCFF009DUY, ENCFF676NPW, ENCFF507MGL, ENCFF379MPS, ENCFF665FHC, ENCFF926FUM, ENCFF113PMT, ENCFF939ZFS, ENCFF927JBT, ENCFF178MOP, ENCFF357NRL, ENCFF176NOI, ENCFF680WBN, ENCFF580QGA, ENCFF917COW, ENCFF657VJX, ENCFF156WRH, ENCFF411DIA, ENCFF967IKR, ENCFF561WNI, ENCFF624NUZ, ENCFF968EAL, ENCFF253FON, ENCFF987VNY, ENCFF485MXM, ENCFF998HOR, ENCFF507XOB, ENCFF334FMW, ENCFF495MHZ, ENCFF465JKF, ENCFF717TWA, ENCFF074EZL, ENCFF410EDG, ENCFF947KPB, ENCFF973LDQ, ENCFF482CEV, ENCFF823SYE, ENCFF478MPX, ENCFF338HQY, ENCFF545DAF, ENCFF036YVI, ENCFF619BDC, ENCFF558DSF, ENCFF716LRI, ENCFF062ARS, ENCFF948TXN, ENCFF990CFV, ENCFF203EDM, ENCFF785ACI, ENCFF423EMU, ENCFF471ZTZ, ENCFF544XKC, ENCFF407AOX, ENCFF182QDI, ENCFF398EQF, ENCFF019PEL, ENCFF641LJY, ENCFF002CXC, ENCFF812QPN, ENCFF019ALY, ENCFF197OGH, ENCFF284LRP, ENCFF944GJH, ENCFF968KBN, ENCFF706LIT, ENCFF249EZR, ENCFF494ZUY, ENCFF296CNQ, ENCFF723RSX, ENCFF440JJW, ENCFF699ZII, ENCFF340PPI, ENCFF475TQE, ENCFF938IOJ, ENCFF058NMZ, ENCFF300XUA, ENCFF083YCQ, ENCFF786WZD, ENCFF872JJJ, ENCFF998YJY, ENCFF237UAN, ENCFF366UBB, ENCFF235ZZP, ENCFF175IIE, ENCFF992QZV, ENCFF638YFC, ENCFF582YPB, ENCFF708GFU, ENCFF998KKJ, ENCFF312RFN, ENCFF985TSH, ENCFF890YEO, ENCFF134NMW, ENCFF849ZUF, ENCFF883WYX, ENCFF030QEM, ENCFF881XQF, ENCFF732HOE, ENCFF144PPR, ENCFF718NUA, ENCFF664XPS, ENCFF966MVC, ENCFF573DRR, ENCFF804ABO, ENCFF868QLL, ENCFF502KHR, ENCFF012VES, ENCFF613RNG, ENCFF114IWY, ENCFF682SIY, ENCFF209MQX, ENCFF456PQY, ENCFF368TYM, ENCFF660VPM, ENCFF429XKT, ENCFF137IBM, ENCFF822FWM, ENCFF858QMI, ENCFF929YBC, ENCFF925DJG, ENCFF577UJR, ENCFF706SJZ, ENCFF096XMD, ENCFF002CXA, ENCFF591BIT, ENCFF394LOQ, ENCFF337DKJ, ENCFF297JZQ, ENCFF255EOB, ENCFF455TDM, ENCFF328XKC, ENCFF821VBM, ENCFF191MPD, ENCFF667HLC, ENCFF048OIZ, ENCFF633VCV, ENCFF242IWJ, ENCFF294EHN, ENCFF386WKY, ENCFF157UUF, ENCFF007DKB, ENCFF820EVZ, ENCFF522JUV, ENCFF511QHY, ENCFF664ZGR, ENCFF911VSD, ENCFF914RFS, ENCFF126BCK, ENCFF712AXK, ENCFF168HAG, ENCFF332ICQ, ENCFF030HWZ, ENCFF490VVG, ENCFF041YQC, ENCFF833FCO, ENCFF150VTD, ENCFF801YZV, ENCFF167HYX, ENCFF179NDS, ENCFF010MMQ, ENCFF517YCC, ENCFF422NGZ, ENCFF838COI, ENCFF253NXR, ENCFF954NAJ, ENCFF002DDJ, ENCFF542DOG, ENCFF986MHU, ENCFF738WCE, ENCFF996IVU, ENCFF558JBX, ENCFF833KXP, ENCFF294VWT, ENCFF080NBZ, ENCFF057ZUY, ENCFF197YHU, ENCFF399GAE, ENCFF809OEX, ENCFF744SVC, ENCFF539WLH, ENCFF666LZV, ENCFF622RBW, ENCFF462FRU, ENCFF085HTY, ENCFF365NKO, ENCFF921WMN, ENCFF883TOD, ENCFF010STZ, ENCFF694BIA, ENCFF120IDE, ENCFF971VJZ, ENCFF175IUO, ENCFF920RYJ, ENCFF294HEI, ENCFF211XFR, ENCFF588OLK, ENCFF937VZI, ENCFF184RRU, ENCFF996CUX, ENCFF617CAZ, ENCFF142CPK, ENCFF644WLI, ENCFF404YMX, ENCFF768TJI, ENCFF320THV, ENCFF872YNU, ENCFF836VRV, ENCFF561USY, ENCFF511OXK, ENCFF034PQD, ENCFF049CRW, ENCFF434QZL, ENCFF396NSO, ENCFF322DSX, ENCFF562JDH, ENCFF360BTO, ENCFF925ANU, ENCFF394IKH, ENCFF484HCG, ENCFF995NIE, ENCFF549KOD, ENCFF785HXY, ENCFF948YVM, ENCFF248IWJ, ENCFF530FZL, ENCFF273TYA, ENCFF738TKN, ENCFF549TYR

**Supplementary Table S4.** Number fragments for all chromHMM (1) states in GM12878 and K562. The detailed listing over all fragments included in chromHMM for the hg19 human genome assembly obtained with the UCSC Table Browser (2).

| | **K562** | **GM12878** |
|---|---|---|
| 1_Active_Promoter | 15625 | 15278 |
| 2_Weak_Promoter | 30008 | 35065 |
| 3_Poised_Promoter | 6266 | 5263 |
| 4_Strong_Enhancer | 30122 | 25486 |
| 5_Strong_Enhancer | 38974 | 38604 |
| 6_Weak_Enhancer | 58856 | 69111 |
| 7_Weak_Enhancer | 121889 | 109468 |
| 8_Insulator | 41147 | 33265 |
| 9_Txn_Transition | 21174 | 16227 |
| 10_Txn_Elongation | 24867 | 26509 |
| 11_Weak_Txn | 100582 | 82312 |
| 12_Repressed | 38902 | 25483 |
| 13_Heterochrom/lo | 78279 | 75112 |
| 14_Repetitive/CNV | 7228 | 8028 |
| 15_Repetitive/CNV | 8338 | 6128 |

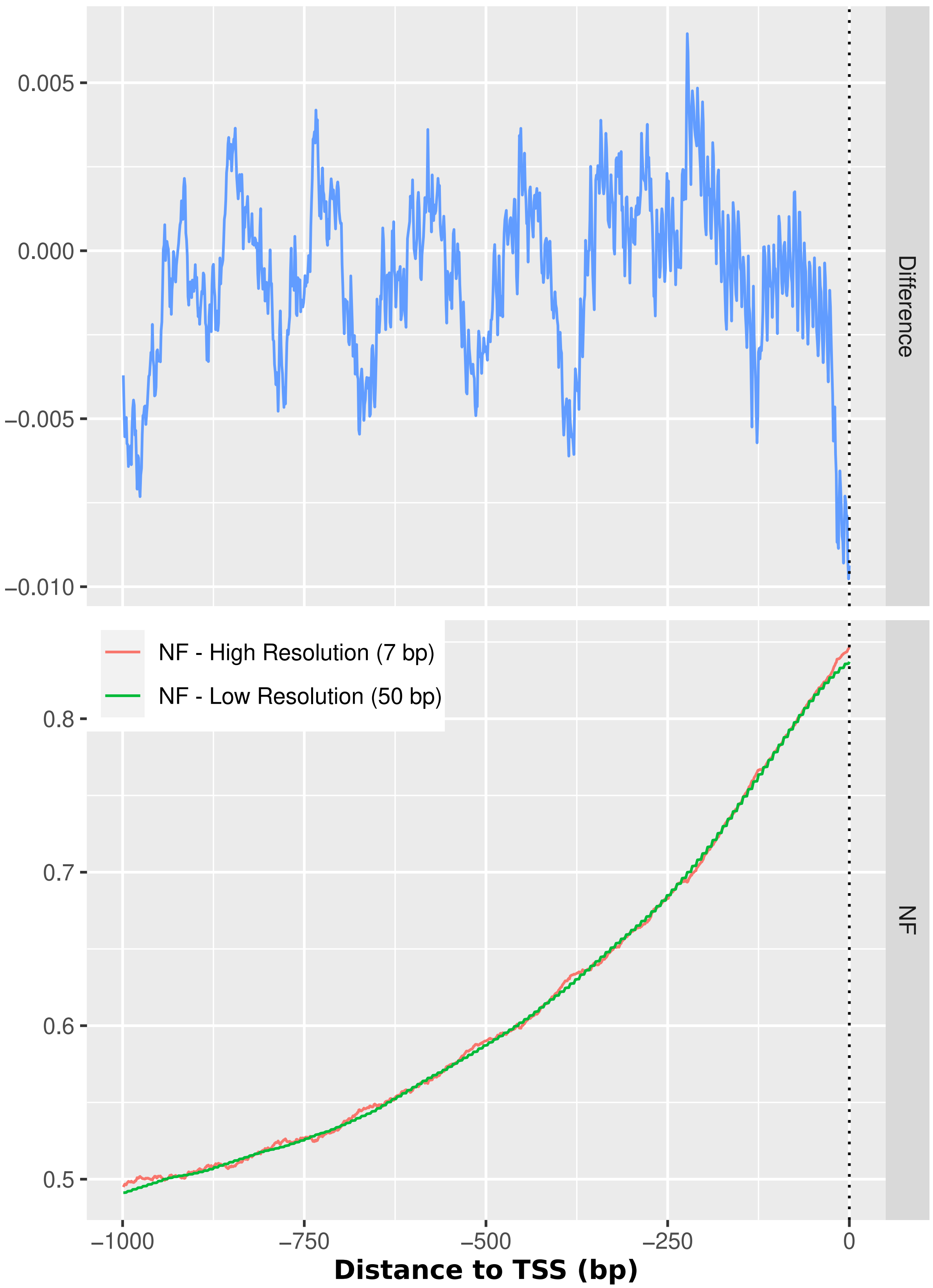


**Supplementary Figure S1. NF score resolution Comparison**. We calculated and provide two resources. The High resolution NF score is calculated for all human promoters and is built by calculating the relative frequency of nucleosomal DNA predictions with the classifier for overlapping sliding windows with a step size of 7 with the result of a score between 0 and 1. In contrast, the low resolution score is created by applying the classifier in a binary way, so that each window yields an array of 50 central bp within a 147 bp window of either 0 (linker) or 1 (nucleosomal). The profiles show the mean score distribution of 10000 randomly sampled promoters from RefSeq. The vertical line at 0 bp denotes the TSS.

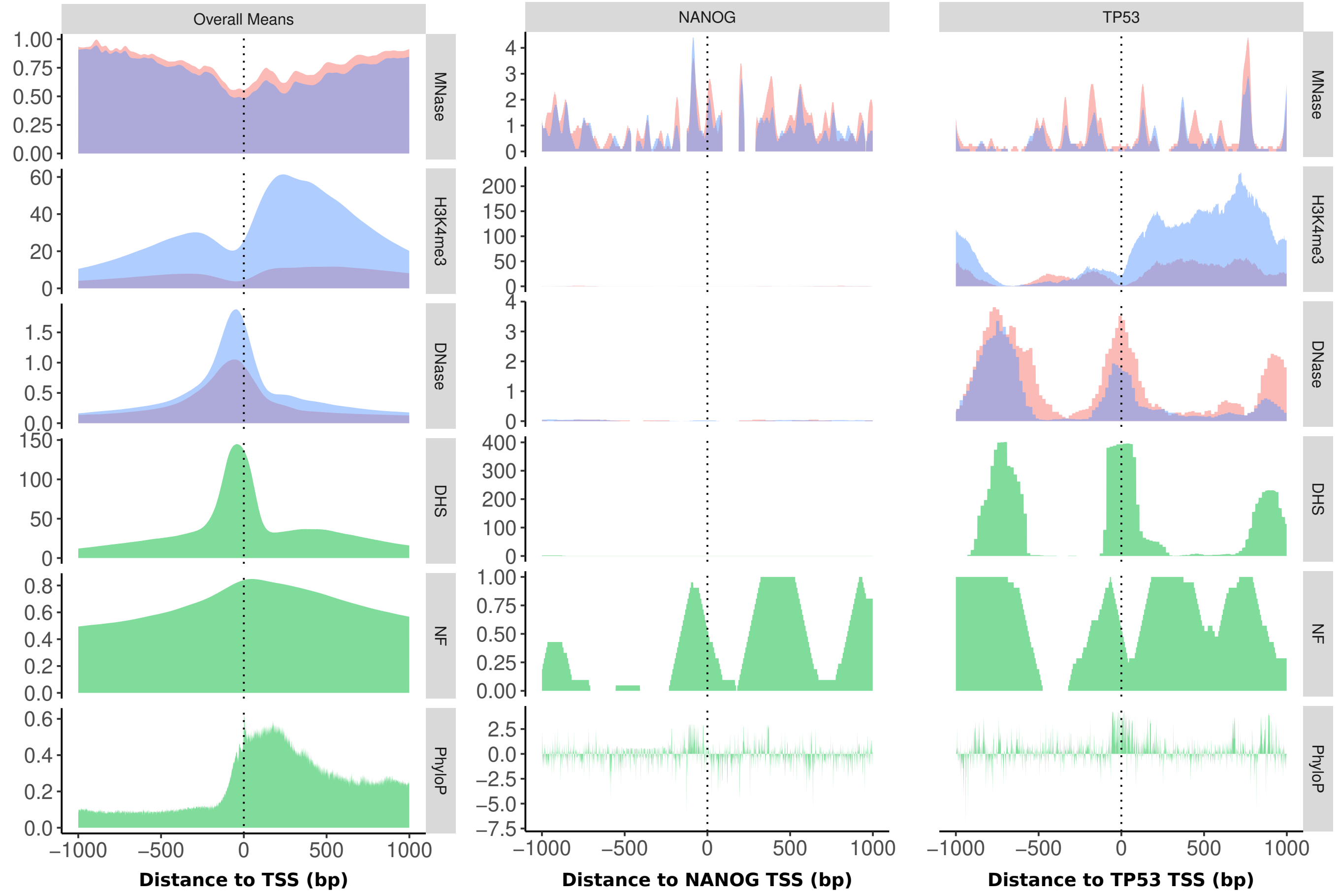


**Supplementary Figure S2. Promoter Profiles**. Profiles for experimental (MNase-seq, DNase-seq, H3K4me3 ChIP-seq) and global data (green: DHS score, NF score, phyloP) as a mean of all human RefSeq promoters and the promoters of the genes NANOG and TP53 (B-C) in K562 (blue) and GM12878 (red). This plot is an extension of Figure 3 in the main paper, showing a larger portion of the promoters downstream of the TSS.

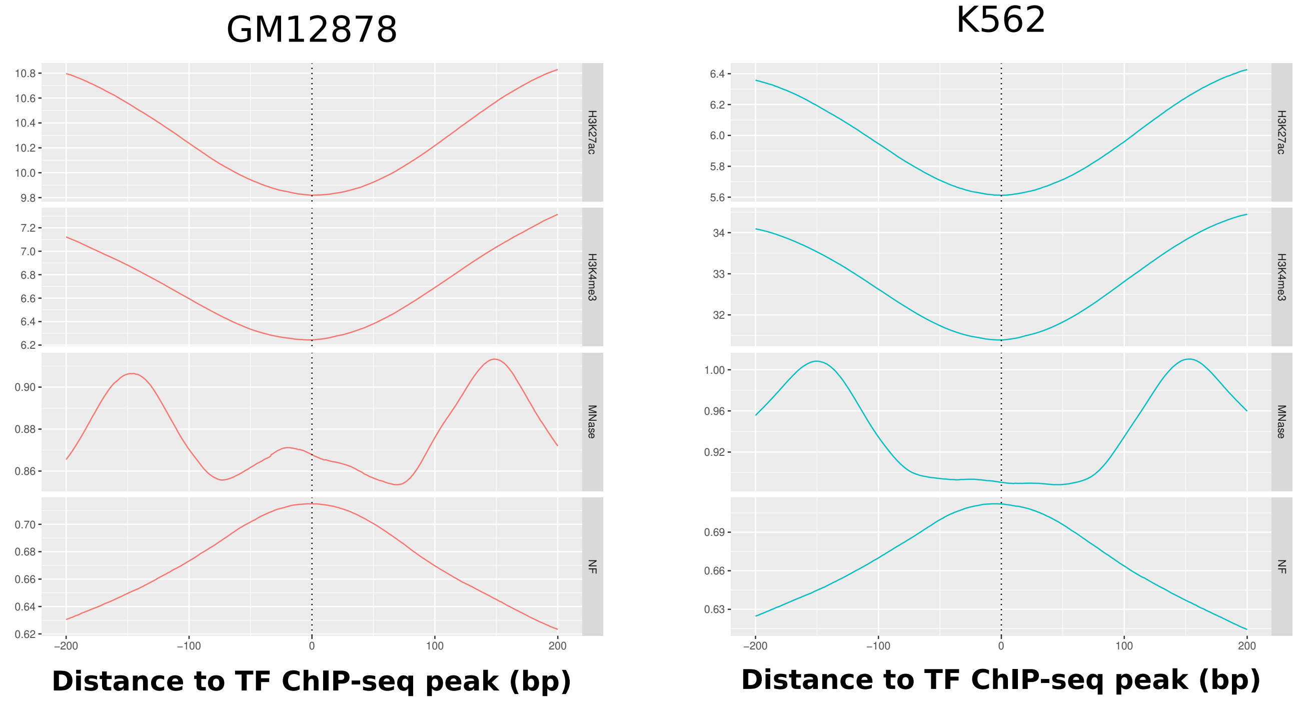


**Supplementary Figure S3. TF Profiles**. Extended figure relating to Figure 4 B in the main manuscript. Sequence-intrinsic Nucleosome support, nucleosome occupancy and histone modification signal for H3K4me3 and H3K27ac around the ChIP-seq peaks of all human TFs from ENCODE in hg19 as a mean profile surrounding the point-source peak. The sequence-intrinsic nucleosome support is highest around the TF binding locations. At the same time the nucleosome occupancy signal is depleted. The profiles for both histone modifications are widely decreased around the ChIP-seq peaks, showing that both promoters and enhancers are expected to behave similarly in regards to the competition between TFs and nucleosome.
*Y-Axes: H3K27ac - ChIP-seq control normalized tag density; H3K4me3 - ChIP-seq control normalized tag density; DNase - read depth normalized signal; MNase - density graph of signal enrichment; NF - NF score.*

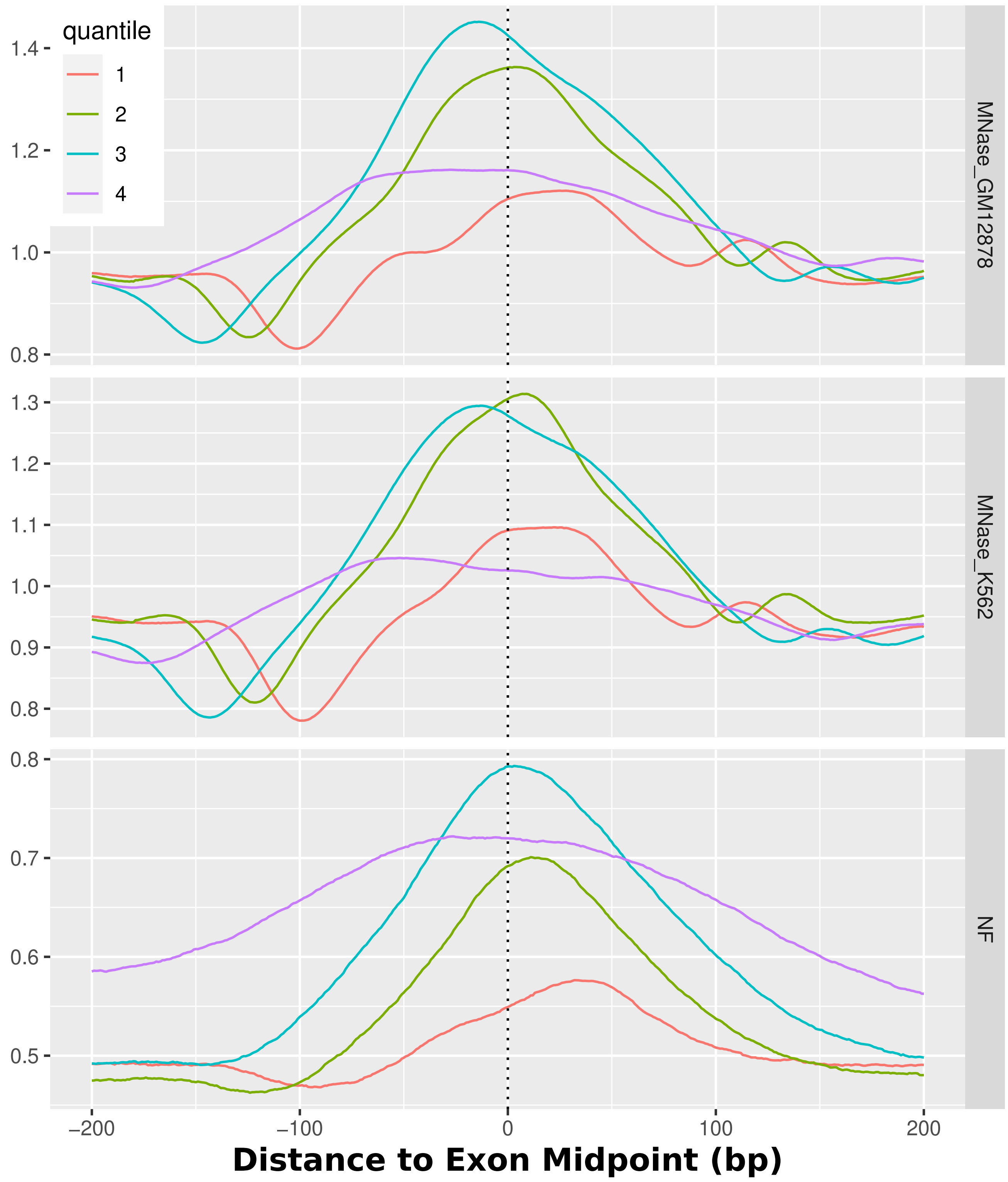


**Supplementary Figure S4. Nucleosomes around Exon Midpoints**. MNase-seq and NF scores for all human exons, centered around the center of all exons. The quantiles separate different lengths of Exons. The lower quantiles denominate shorter exons. In particular, they denominate these exon lengths: 1 = 1-90 bp (median 69 bp), 2 = 90-129 bp (median 109 bp), 3 = 129-190 bp (median 153 bp), 4 = 190-91671 bp (median 333 bp). The strand orientation is taken into account and always 5'-3' directed. The NF score shows a high sequence-defined nucleosome support for exon regions and experimentally a high nucleosome occupancy can be observed. The longer exons generally display a higher NF score.

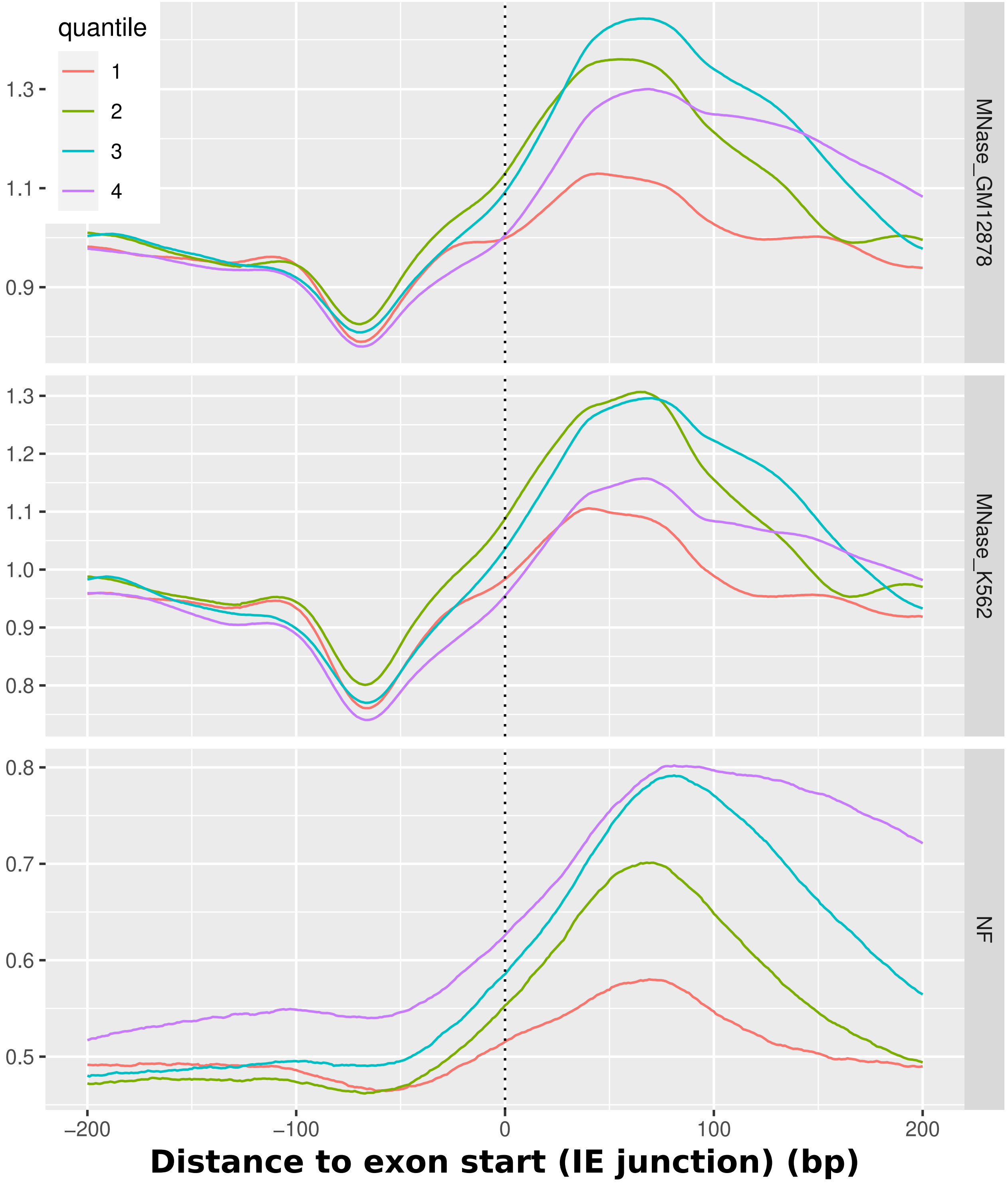


**Supplementary Figure S5. Nucleosomes around Exon Starts**. MNase-seq and NF scores for all human exons, centered around the Intron/Exon junctions. The quantiles separate different lengths of Exons. The lower quantiles denominate shorter exons. In particular, they denominate these exon lengths: 1 = 1-90 bp (median 69 bp), 2 = 90-129 bp (median 109 bp), 3 = 129-190 bp (median 153 bp), 4 = 190-91671 bp (median 333 bp). The strand orientation is taken into account and always 5'-3' directed. The nucleosome occupancy is highest downstream of the TSS for all exon lengths. The IE junction is based on the apex of the rise of nucleosome positioning from an upstream depletion, around 75 bp in distance, towards a clearly positioned nucleosome 75 bp downstream.

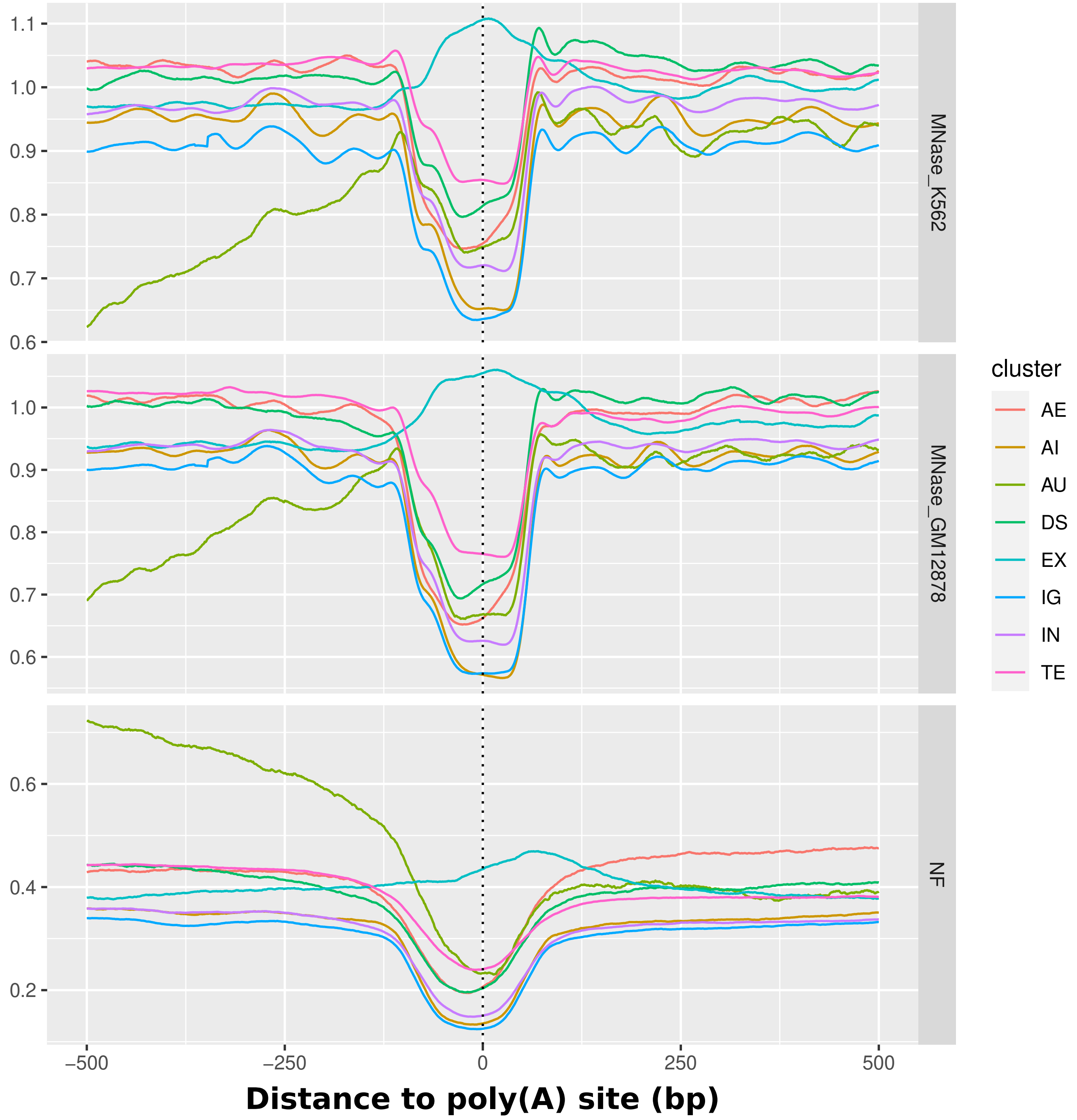


**Supplementary Figure S6. Poly(A) Sites**. Detailed overview over all poly(A) sites from the PolyASite 2.0 atlas, separated by cluster (AE = Anti-sense to an exon, AI = Anti-sense to an intron, AU = 1000 nt upstream in anti-sense direction of a transcription start site, DS = 1000 nt downstream of an annotated terminal exon, EX = Exonic, IG = Intergenic, IN = Intronic), TE= Terminal Exons (3). All groups apart from exonic Poly(A) sites exhibit a local decrease around the site both in sequence-defined nucleosome support (nf) and nucleosome occupancy in both cell lines K562 and GM12878.

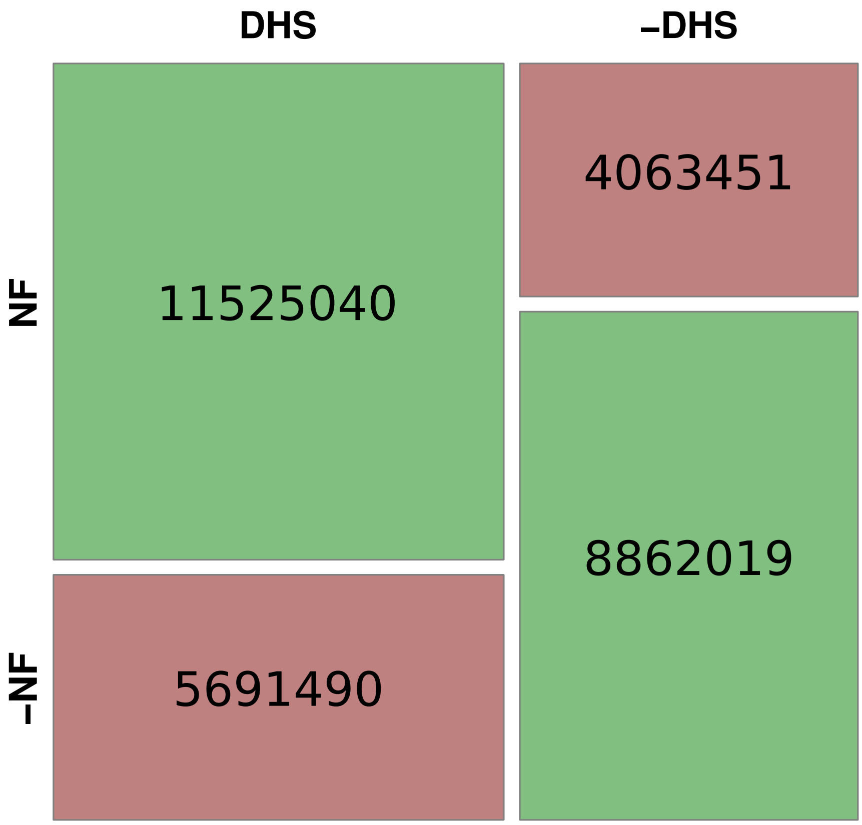


**Supplementary Figure S7. Sequence-Defined**. In order to show the relation between highly important gene regulatory regions and the sequence-intrinsic nucleosome support, we compared the NF score and the DHS score (see Methods section) with the Fisher test. For that purpose, we took the individual base pairs of all human promoters and divided both measures into two bins each, according to their median. The median for the high-resolution NF score is at 0.76 and the DHS score at 1 (i.e. 1 out of 403 primary cell lines is open at that particular bp). There is a significant (Fisher test, p-value $<$2.2e-16, odds ratio: 4.416, 95% confidence interval: [4.409 4.423] ) relation between regions with a high sequence-intrinsic nucleosome support and accessible chromatin *in vivo*, indicating a co-evolution of nucleosome as well as transcription factor attraction in human promoters.

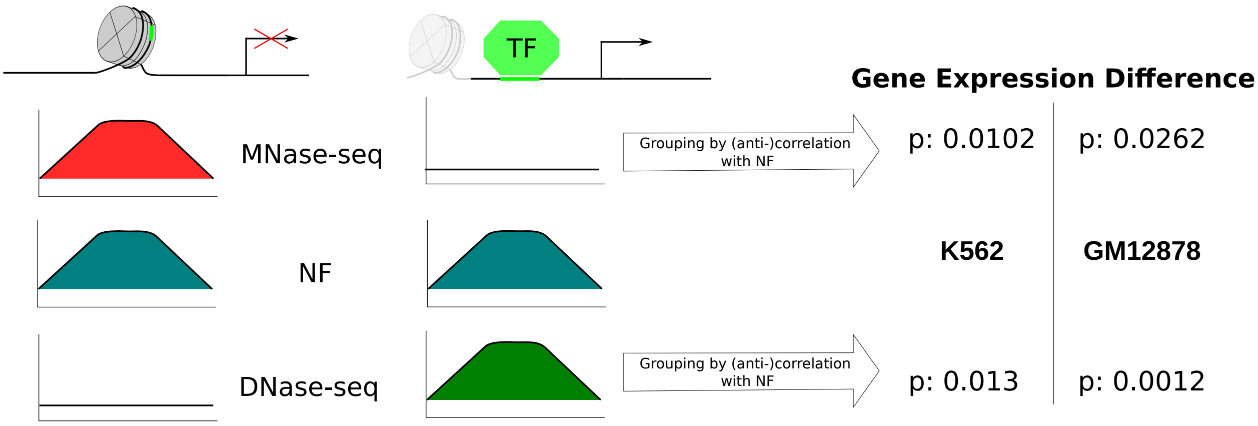


**Supplementary Figure S8. Sequence-positioned nucleosomes affect gene expression**. We calculated the Spearman correlation between the high-resolution NF score in all human promoters with the respective MNase-seq and DNase-seq signal per promoter. Then we tested whether the gene expression is lower when NF score and MNase-seq show a significant positive correlation vs. a negative correlation. Respectively, we tested for a higher gene expression in genes with a positive correlation of NF score and DNase-seq vs. gene expression in negatively correlated genes. The table shows the p-values of the wilcoxon test per cell line (K562, GM12878) and comparison (MNase-seq, test direction "less"; DNase-seq, test direction "greater"). The p-values are adjusted with the Bonferroni-Holm method. The significant p-values suggest that there is an influence on gene expression based on the displacement of sequence-intrinsically positioned, "regulatory" nucleosomes. The definition of promoters follows the methods of the main paper. The gene expression values are obtained from ENCODE (GM12878: ENCFF873VWU (rep1), ENCFF345SHY (rep2); K562: ENCFF928NYA (rep1), ENCFF003XKT (rep2)) (4)

.
